# Divergent chromatin remodeling in post-sepsis MDSCs underlies MHC class II repression in CCI

**DOI:** 10.64898/2026.01.21.700934

**Authors:** Jason O. Brant, Marie-Pierre L. Gauthier, Christine E. Rodhouse, Ruoxuan Wu, Miguel Hernández-Ríos, Leilani Zeumer-Spataro, Marvin L. Dirain, Ricardo F. Ungaro, Ivanna L. Rocha, Whittman B. Wiggins, Angel M. Charles, Feifei Xiao, Letitia E. Bible, Alicia M. Mohr, Shawn D. Larson, Jaimar C. Rincon, Shannon M. Wallet, Maigan A. Brusko, Tyler J. Loftus, Lyle L. Moldawer, Clayton E. Mathews, Rhonda Bacher, Guoshuai Cai, Robert Maile, Philip A. Efron, Michael P. Kladde

## Abstract

Sepsis survivors exhibit divergent clinical trajectories, including rapid recovery (RAP) or progression to chronic critical illness (CCI), yet how these outcomes are linked to epigenetic repression remains poorly defined. Here, we applied an optimized Omni-ATAC approach to profile chromatin accessibility in CD66b⁺ myeloid-derived suppressor cells (MDSCs) from healthy participants and longitudinally sampled sepsis cohorts stratified by outcome. RAP samples progressively restored healthy chromatin states, whereas CCI samples remained epigenetically fixed in aberrant configurations. Chromatin remodeling exhibited strong pathway specificity: promoters associated with MHC class II antigen presentation were coordinately repressed in CCI, while MHC class I antigen processing and presentation machinery remained preserved. Genome-wide analyses revealed extensive promoter remodeling during recovery in RAP, including immune regulatory loci such as *ARG1*, *CD274*, and *S100A8*/*A9*, contrasted with broad suppression of immune, metabolic, and chromatin regulatory programs in CCI. These findings define divergent epigenetic trajectories in post-sepsis MDSCs and implicate selective failure of antigen presentation as a mechanism of sepsis-induced immunoparalysis in CCI.

## Introduction

Sepsis, defined as life-threatening organ dysfunction caused by dysregulated host response to infection (Singer *et al*, 2016), is among the most morbid conditions encountered in clinical practice (Rudd *et al*, 2020). Recognized by the World Health Organization as a global health threat (Global Sepsis Alliance, n.d.), sepsis remains the most resource-intensive cause of inpatient hospitalization in the United States (Paoli *et al*, 2018). Although advances in early recognition and acute management have reduced in-hospital mortality (Kim *et al*, 2024), long-term morbidity and mortality remain high. This burden is expected to increase as sepsis incidence rises in an aging population with increasing chronic comorbidities (Shankar-Hari & Rubenfeld, 2016).

Survivors of sepsis typically experience one of two clinical outcomes: 1) rapid recovery (RAP) or 2) progression to chronic critical illness (CCI) (Darden *et al*, 2021b). CCI is defined as a prolonged ICU requirement (>14 days) with persistent organ injury and dysfunction (Brakenridge *et al*, 2019; Darden *et al*., 2021b) and is further characterized by recurrent infections and immune dysregulation, along with impaired functional recovery and increased late mortality, disproportionately affecting older males (Carson, 2012; Kahn *et al*, 2015; Nelson *et al*, 2010).

Our work and that of others indicate that CCI patients develop pathologic myeloid differentiation contributing to Persistent Inflammation, Immunosuppression, and Catabolism Syndrome (PICS) (Darden *et al*, 2021c). This dysfunctional differentiation of immature myeloid cells includes an overabundance of immunosuppressive myeloid-derived suppressor cells (MDSCs) in CCI sepsis survivors (Mathias *et al*, 2017a). First discovered in cancer, MDSCs can promote both chronic low-grade inflammation and immunosuppression of host leukocytes (e.g., lymphocytes) (Talmadge & Gabrilovich, 2013). We have demonstrated that MDSCs arising after sepsis are associated with CCI and exhibit unique metabolic and immune phenotypes (Barrios *et al*, 2024). Although FDA-approved immunomodulators can alter MDSC function in oncologic settings, our work demonstrates that MDSCs from sepsis patients operate through mechanisms and functional programs distinct from those of cancer-associated MDSCs (Barrios *et al*., 2024). Thus, precision medicine approaches will be required to modify these cells in sepsis patients, either in circulation or during their generation or differentiation from hematopoietic stem cells.

Epigenetic mechanisms, including nucleosome positioning, histone modifications, and DNA methylation, govern the generation of stable programs of tissue- and cell-specific expression during differentiation (Zentner & Henikoff, 2013). Environmental stimuli also elicit dynamic reprogramming of enhancer and promoter chromatin, which typically reverses after stimulus resolution (Minnoye *et al*, 2021). Establishing both stable and reversible epigenetic states requires modulation of chromatin accessibility, largely dictated through remodeling of nucleosome occupancy or positioning, particularly at transcription start sites (TSSs). These accessibility changes are more closely tied to cellular identity and specification than transcriptomic profiles (Klemm *et al*, 2019). Genome-wide changes in chromatin accessibility can be profiled using Omni-ATAC, an improved Assay for Transposase-Accessible Chromatin using sequencing (ATAC-seq) (Corces *et al*, 2017).

Given our previous transcriptomic results (Barrios *et al*., 2024), we hypothesized that CD66b^+^ MDSCs isolated from peripheral blood mononuclear cells (PBMCs) of sepsis survivors would exhibit significant epigenetic differences across cohorts defined by time after sepsis diagnosis and clinical outcome (RAP versus CCI). To test this, we applied the Omni-ATAC protocol further optimized for CD66b^+^ MDSCs, enabling robust profiling of chromatin accessibility in this challenging cell population. Using this approach, we uncovered distinct, outcome-associated chromatin accessibility patterns distinguishing early-death (mortality within 14 days of sepsis diagnosis) and CCI cohorts from RAP. PBMC CD66b^+^ MDSCs from all septic patients showed substantial differences in global chromatin accessibility relative to HP-derived samples. Furthermore, over time, chromatin accessibility patterns in RAP patients progressively regained baseline chromatin states, whereas CCI patients remained epigenetically ‘locked’ in aberrant states. We further hypothesized that these divergent epigenetic trajectories would extend to key immunoregulatory pathways governing antigen presentation. Notably, we found pathway-specific alterations in chromatin accessibility differentially impacted MHC class II and MHC class I gene programs, thereby contributing to impaired adaptive immune function in CCI.

## Results

### PBMC CD66b^+^ immunosuppression differs by clinical status and sex

Co-culture of PBMC CD66b^+^ cells derived from CCI patients at day 14 (d14) post-sepsis with total CD3^+^ T lymphocytes significantly reduced proliferation of CD8^+^ T cells (Supplementary Fig. 1A), but not the CD4^+^ T-cell subpopulation (Supplementary Fig. 1B). In contrast, CD66b^+^ cells from RAP survivors at d14 (Supplementary Fig. 1A, B), as well as from both RAP and CCI cohorts at d4 post-sepsis (data not shown), did not elicit this effect. Notably, at 6 months (6mo) post-sepsis, the fold change in proliferation of stimulated CD8^+^ and CD4^+^ T lymphocytes was reduced to a greater extent by addition of CD66b^+^ cells derived from CCI compared with RAP survivors, indicating more pronounced T-cell suppression by CCI-derived cells (Supplementary Fig. 1C, D). Together, these data confirm that PBMC CD66b^+^ cells isolated at d14 meet functional criteria for MDSCs.

### Distinct chromatin accessibility trajectories distinguish sepsis recovery status

We next asked whether functional differences in PBMC CD66b^+^ cell activity were associated with distinct chromatin accessibility profiles. To address this question, we applied a further optimized Omni-ATAC protocol to circulating MDSCs (PBMC-derived CD66b^+^ cells) from healthy participants (HPs) and sepsis patients stratified by clinical status: RAP, CCI, and early death (mortality at <14 days, before clinical status could be determined). The cohort included both males (n = 20) and females (n = 9) subjects. For sepsis patients, samples were collected at multiple time points—day 4 (d4), day 14-21 (d14), and 6 months (6mo) post-sepsis onset—yielding a total of *N* = 48 samples.

Dimensionality reduction analyses, including principal component analysis (PCA) and uniform manifold approximation and projection (UMAP), revealed distinct clustering of samples by clinical status and time point (Fig. 1). To minimize sex-specific effects on chromatin accessibility, only autosomal data were used, and sex was included as a covariate during batch correction when male and female samples were analyzed together. HP samples clustered together and were clearly separated from sepsis samples along the first two dimensions. Early RAP samples (d4) showed the greatest deviation from HP and clustered near CCI samples. Both RAP and CCI samples exhibited temporal trajectories in chromatin accessibility space, with d14 and 6mo samples shifting toward the HP cluster, consistent with recovery (Fig. 1A; Fig. 1B, top panel). However, except for one patient, CCI sample trajectories were less pronounced than those of RAP, with most CCI samples shifting only a short distance and seldom crossing dimension axes, potentially reflecting delayed or incomplete recovery (Fig. 1A; Fig. 1B, bottom panel). Group-level clustering was not driven by batch effects, as PCA colored by Omni-ATAC processing batch shows no batch-associated clustering (Supplementary Fig. 2A).

**Figure 1.**
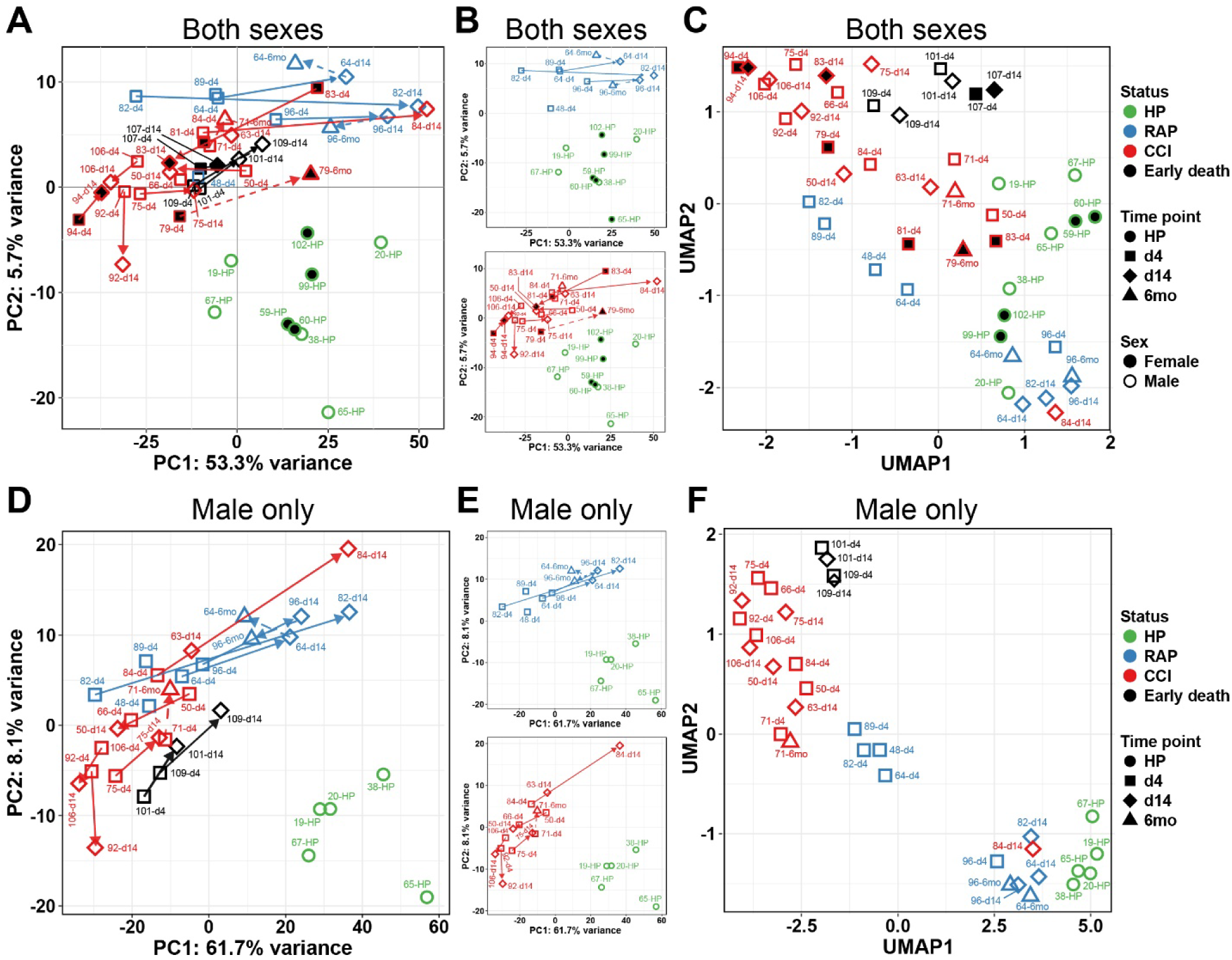
Differential accessibiliy analysis identifies dynamic chromatin remodeling in RAP recovery versus CCI. **A** PCA of top 1,000 most-variable peaks. **B** Same data as in panel **A**, but with samples removed to focus specifically on RAP (top panel) or CCI (bottom panel) clinical groups. Points are colored by clinical status (HP, RAP, CCI, Early death) and shaped by time point; sex is indicated by filled shapes for Female, no fill for Male. Text labels are sample IDs; arrows show within-patient trajectories: solid arrow = d4 → d14, dashed arrow = d14 → 6mo, dashed direct arrow = d4 → 6mo when d14 is missing. Arrow direction indicates the temporal progression of each patient in PCA space; arrow length is the Euclidean displacement of the projected samples. Group sizes: HP (n = 11), RAP (n = 10), CCI (n = 21), Early death (n = 6). **C** UMAP (n_neighbors = 5, min_dist = 0.01; input = top 500 most-variable peaks, PCs = 30). **D** PCA of male-only subset; status and time points labeled as in panel **A**. **E** Same data as in panel **D**, but simplified as in panels **B**. **F** UMAP (n_neighbors = 4, min_dist = 0.01; input = top 500 most-variable peaks, PCs = 30) of male-only subset; status and time point labeled as in panel **A.** Male only group sizes: HP (n = 6), RAP (n = 10), CCI (n = 14), Early death (n = 4).

Because PCA primarily captures global variance, we also generated UMAP to visualize local data structure. UMAP revealed clustering patterns by clinical status and time post-sepsis that were similar to those observed by PCA (Fig. 1C). Analysis of male-only samples (*n* = 34) showed clearer separation (PCA and UMAP) and trajectory structure (PCA) (Fig. 1D, F). The female-only subset (*n* = 14), which lacked RAP-derived samples, nonetheless exhibited distinct clustering of samples from HP, CCI, and early-death patients (Supplementary Fig. 2B, C).

### Differential chromatin accessibility reveals extensive remodeling across clinical states and time

Differential accessibility analysis was performed for pairwise comparisons between condition groups (n = 48). Eight contrasts were evaluated; differentially accessible regions (DARs) were considered significant if the global adjusted *P* value was ≤ 0.01 and the absolute log_2_ fold change (log_2_(FC)) exceeded 1 (≥ 2-fold). The number of DARs varied markedly by comparison, with the largest numbers observed in RAP d14 samples and in longitudinal comparisons across time (RAP d14 versus RAP d4) (Table 1; Supplementary Table 1 lists annotated DARs for each contrast).

Compared with HPs, both RAP and CCI subjects displayed modest changes on d4 (CCI: 104 DARs, 53 increased and 51 decreased accessibility; RAP: 42 DARs, 8 increased and 34 decreased accessibility) (Fig. 2A, B; Table 1). Whereas the number of DARs in CCI subjects remained limited at d14 (72 DARs, 65 increased and 7 decreased accessibility), RAP subjects exhibited dramatic accessibility changes by d14 (3,900 DARs: 3,897 increased and 3 decreased accessibility) (Fig. 2C, D; Table 1).

**Figure 2.**
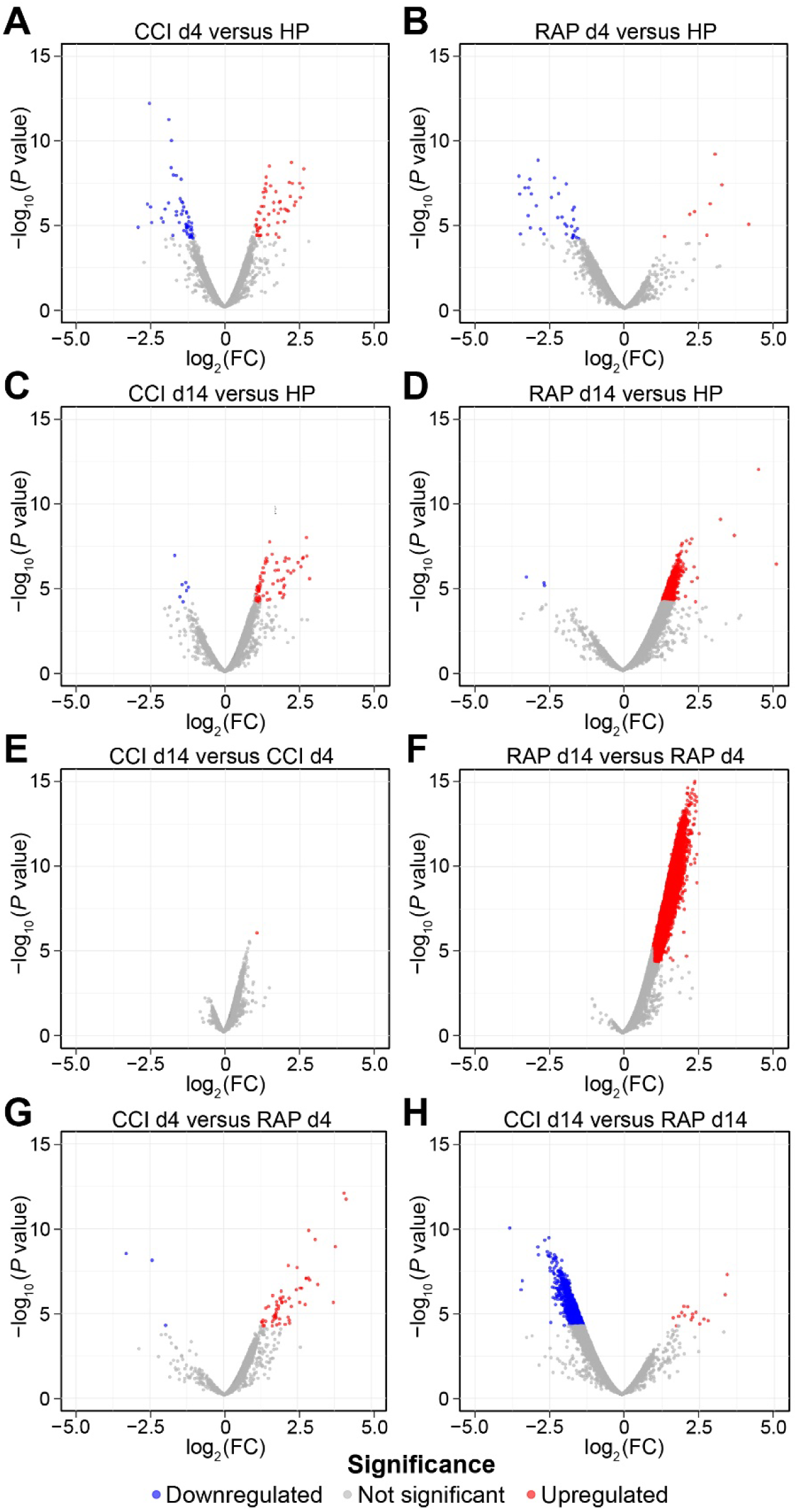
Volcano plots of differential chromatin accessibility comparing clinical status and time point within and across groups. RAP and CCI clinical groups at d4 and d14 versus HP (**A-D**), temporal changes within CCI and RAP across time (**E** and **F**), and between-group comparisons for CCI and RAP at d4 and d14 (**G** and **H**). Each DAR is plotted by the -log_10_ of raw *P* value on y-axis and the log_2_(Fold Change, FC) on the x-axis. DARs were considered significant if the global adjusted *P* value was ≤ 0.01 and the log_2_(FC) was ≥ 1.0. Significant DARs are colored blue for decreased accessibility and red for increased accessibility. Gray indicates no significant change in accessibility.

Temporal changes between d4 and d14 in RAP patients showed extensive chromatin remodeling (22,970 DARs, all showing increased accessibility at d14) (Fig. 2E, F; Table 1). Direct comparisons between clinical states revealed relatively few differences at d4 (61 DARs between CCI and RAP, 58 of which increased in CCI) (Fig. 2G; Table 1). By contrast, there was substantial divergence by d14 (2,905 DARs), with CCI predominantly showing decreased accessibility relative to RAP (2,889 decreased versus 16 increased DARs) (Fig. 2G, H; Table 1).

Interpreting the biological meaning of DARs outside of regulatory elements is challenging without *a priori* knowledge of their function. Therefore, we next focused on the subset of DARs that mapped to promoter regions (±1 kb of TSSs). As with total DARs, RAP and CCI patients at d4 had only minimal promoter DARs compared with HPs (Supplementary Fig. 3A, B). By d14, RAP subjects exhibited pronounced promoter remodeling (630 DARs, 627 with increased accessibility) (Supplementary Fig. 3C, D). In contrast, CCI patients again showed minimal change across time, whereas RAP patients displayed extensive promoter remodeling between d4 and d14 (13,271 DARs, all with increased accessibility), indicating widespread regulatory chromatin remodeling during rapid recovery (Supplementary Fig. 3E, F). Direct comparisons between conditions revealed few promoter differences at d4 (6 DARs between CCI and RAP), but a striking divergence by d14 (905 DARs), with CCI showing predominantly decreased promoter accessibility relative to RAP (889 decreased DARs) (Supplementary Fig. 3G,H). This points to fundamentally different regulatory programs between these conditions at later times when RAP patients are recovering.

UpSet plots were generated to define unique and intersecting genes associated with promoter DARs across clinically relevant comparisons (Supplementary Fig. 4A). Of the 840 genes with promoter DARs distinguishing CCI and RAP at d14, 829 were also present in RAP d14 versus HP and the RAP d14 versus RAP d4 comparisons. In contrast, of the 7,304 genes with promoter DARs in RAP d14 versus RAP d4, more than 80% (6,117) were unique to this contrast and not observed in comparisons with HP or CCI at d14 (Supplementary Fig. 4A), suggesting that longitudinal remodeling during rapid recovery represents a distinct regulatory program rather than a general response to sepsis.

To address the specificity of the pronounced increase in chromatin accessibility observed in RAP d14 promoters, we examined accessibility patterns across two datasets: all ATAC-seq peaks and promoter-specific peaks. If the increased accessibility in RAP d14 was general, we would expect to observe effects of similar magnitude across both global and promoter DARs. However, effect size analysis revealed a striking divergence between these two datasets. Globally, RAP patients showed modest recovery from d4 to d14 (Cohen’s *d* = 0.32), with d14 accessibility remaining lower than that of HPs (*d* = 0.14 versus HP) (Supplementary Fig. 5; Supplementary Table 2). In sharp contrast, at promoter regions, RAP d14 samples exhibited the strongest effect observed in our analysis (*d* = 0.75 versus RAP d4), exceeding HP levels. This selective increase in accessibility at promoter regions, the primary sites of transcriptional regulation, strongly indicates that increased accessibility of RAP d14-derived MDSCs represents targeted biological compensation rather than a systemic technical issue (Supplementary Fig. 5; Supplementary Table 2). These findings support the hypothesis that RAP patients undergo focused chromatin remodeling at regulatory elements during recovery, distinct from the sustained accessibility suppression observed in CCI patients.

### GO enrichment reveals widespread chromatin remodeling in rapid recovery versus CCI

Gene ontology (GO) enrichment analysis was performed using promoter DARs from CCI versus RAP at d14 and within RAP (d14 versus d4) (Supplementary Table 3). As differential accessibility analyses revealed widespread changes in chromatin accessibility, particularly in RAP patients, we focused on selected terms involved in chromosomal organization and chromatin structure. The dot plot for CCI versus RAP at d14 shows that terms related to nucleosome organization, histone methyltransferase complexes, and SWI/SNF activity are enriched among peaks with decreased accessibility. These terms also have smaller gene ratios and higher adjusted *P* values, consistent with broadly repressive chromatin states (Fig. 3A). Connected network (cnet) analysis reinforces this observation, with GO terms related to chromatin remodeling showing exclusively negative log fold changes (Fig. 3B).

**Figure 3.**
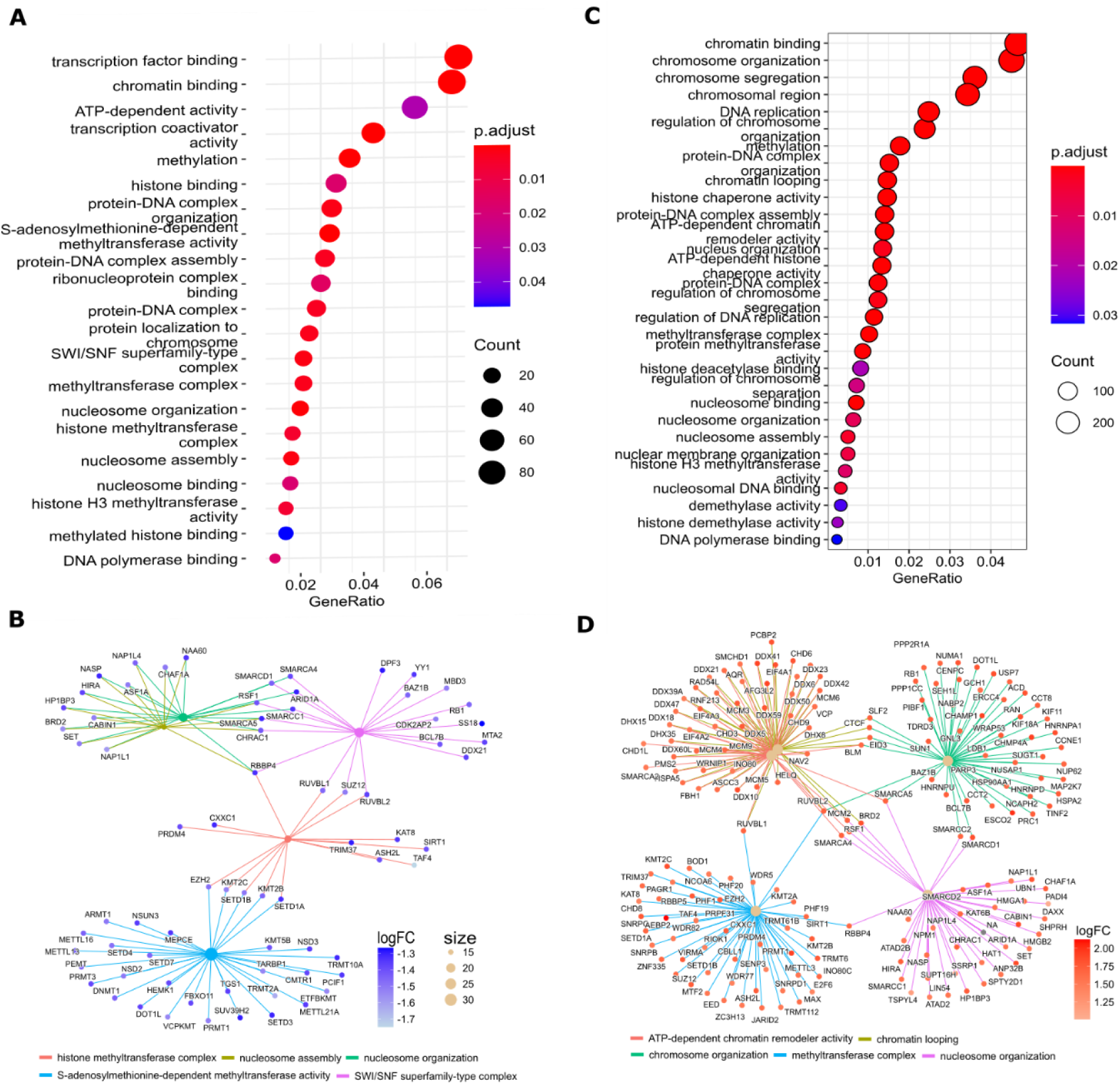
Disruption of chromatin structure and organization in sepsis. **A** Dot plot of selected enriched GO terms from differentially accessible promoters between CCI and RAP at d14. Y-axis is enriched GO term, x-axis is gene ratio (ratio of affected genes to number of genes in pathway), size of the dot represents the number of affected genes per GO term, colors of the dots represent the -log_10_ of the adjusted *P* value (p.adj) of enrichment. **B** Connected network (cnet) plot highlighting selected enriched GO terms related to chromosomal organization and chromatin structure. Central nodes represent enriched GO term categories. Gene nodes are the genes with promoter DARs between CCI vs RAP at d14. Colors of dots associated with each gene represent the log_2_ fold change (logFC) in chromatin accessibility. **C** Dot plot of selected enriched GO terms from promoter DARs from RAP d14 to d4 contrasts. Plot labeled as in panel **A**. **D** Cnet plot of selected enriched GO terms related to chromosomal organization chromatin structure for within RAP comparisons (d14 versus d4), labeled as in panel **B**, except number of genes portrayed limited to the top 50 by enrichment *P* value.

In contrast, the within-RAP temporal comparison (d14 versus d4) shows strong enrichment for chromatin binding, ATP-dependent chromatin remodeler activity, nucleosome assembly, and chromatin organization, reflected by larger gene ratios and lower adjusted *P* values (Fig. 3C). This pattern is further illustrated by the cnet plot of chromatin remodeling modules with positive log fold changes and numerous gene nodes (top 50 shown for clarity) (Fig. 3D).

### IPA identifies pathway-level divergence by clinical status and time

Ingenuity Pathway Analysis (IPA) of promoter DARs revealed a predominantly downregulated pathway profile in MDSCs from CCI patients. Among enriched canonical pathways, most showed negative activation *Z* scores in CCI versus RAP at d14 after sepsis (Fig. 4; Supplementary Table 4), indicating that these pathways are predicted to be functionally inhibited or less active in the CCI group. This widespread pattern of negative *Z* scores indicates global downregulation of immune and metabolic programs in CCI, consistent with an epigenetically repressed state. For example, pathways involved in cytokine signaling (e.g., IL-1, IL-2/IL-7 signaling axes), innate receptor activation (TLR3 signaling), and neutrophil functionality (fMLP signaling and neutrophil degranulation) are all predicted to be inactivated.

**Figure 4.**
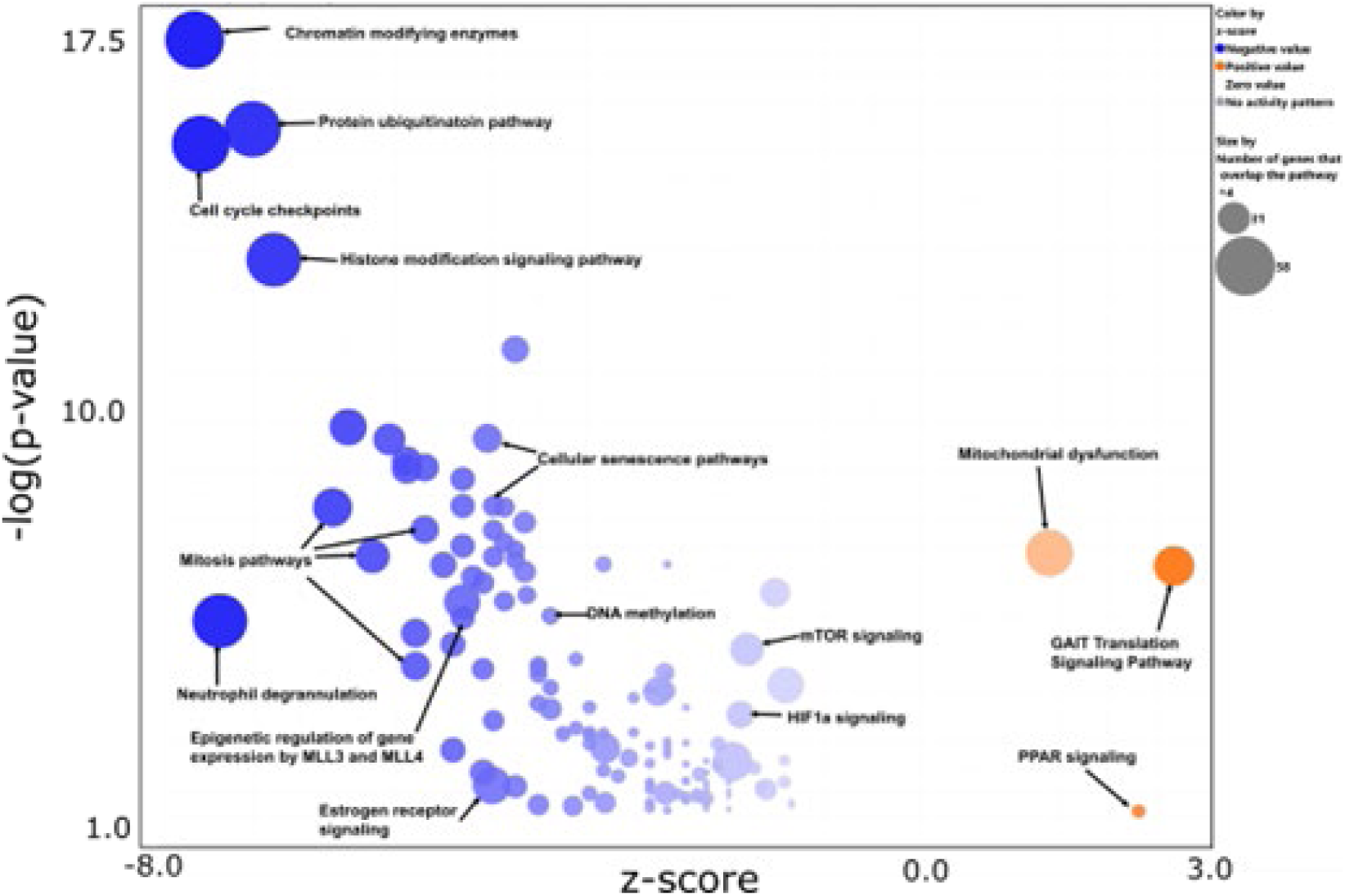
Canonical pathway analysis of differentially accessible promoter regions in CCI versus RAP at d14. Bubble plot of Ingenuity Pathway Analysis (IPA) results from genes with significantly differentially accessible promoter peaks (*P* < 0.01 and |log₂FC| ≥ 1) comparing CCI to RAP at d14 post-sepsis. The x-axis indicates the activation *Z* score, where negative values (blue) represent predicted pathway inhibition in CCI relative to RAP, and positive values (orange) represent predicted activation. The y-axis shows the pathway enrichment significance (–log_10_ of the *P* value). Point size reflects the number of input genes overlapping the given pathway. The majority of enriched pathways in CCI show negative *Z* scores, indicating broad predicted suppression of immune and metabolic programs, while only three pathways show activation (GAIT Translation Signaling, PPAR signaling, and Mitochondrial dysfunction).

Among the most significantly downregulated pathways was Chromatin Modifying Enzymes, which exhibited the strongest negative *Z* score and highest enrichment significance across all canonical pathways analyzed. In contrast, only three pathways exhibited positive *Z* scores (significantly upregulated in CCI): GAIT Translation Signaling, PPAR Signaling, and Mitochondrial Dysfunction. GAIT Translation Signaling (IFN-γ-activated inhibitor of translation) had the highest activation *Z* score (∼2.7), suggesting enhanced activity of translational suppression mechanisms in CCI. PPAR signaling (*Z* score ∼2.3) was also predicted to be activated, reflecting a shift toward lipid metabolic and anti-inflammatory transcriptional programs in MDSCs from CCI patients. In addition, a modest positive *Z* score (∼1.4) for Mitochondrial Dysfunction indicates enrichment of genes associated with impaired mitochondrial function, including the rate-limiting enzyme OGDH and subunits of all five oxidative phosphorylation complexes. Collectively, these findings point to increased mitochondrial stress or inefficiency in CCI relative to RAP. Aside from these three exceptions, however, CCI patient-derived MDSCs demonstrated consistently negative *Z* scores across dozens of pathways, highlighting a broadly suppressed immune-metabolic landscape compared with RAP.

To illustrate chromatin remodeling events underlying these pathway-level changes, we next examined representative genes previously identified in the cancer literature as key mediators of MDSC-induced immunosuppression (Akkari *et al*, 2024). These genes were enriched in multiple GO and IPA pathways in our dataset and exhibited differential promoter accessibility based on time and clinical status (RAP versus CCI), including *ARG1* (encoding arginase), *S100A8* and *S100A9* (encoding alarmins), and *CD274* (encoding PD-L1).

In CD66b^+^ MDSCs from RAP patients, *ARG1* showed a trend toward increased chromatin accessibility at its TSS at d14 compared with HP (Fig. 5A), whereas accessibility at *S100A8*, *S100A9*, and *CD274* promoters increased significantly (Fig. 5B-D; Supplementary Table 5). By 6 months post-sepsis, accessibility at all four promoters in RAP returned to levels comparable to HP, indicating that chromatin remodeling in RAP is dynamic and reversible.

**Figure 5.**
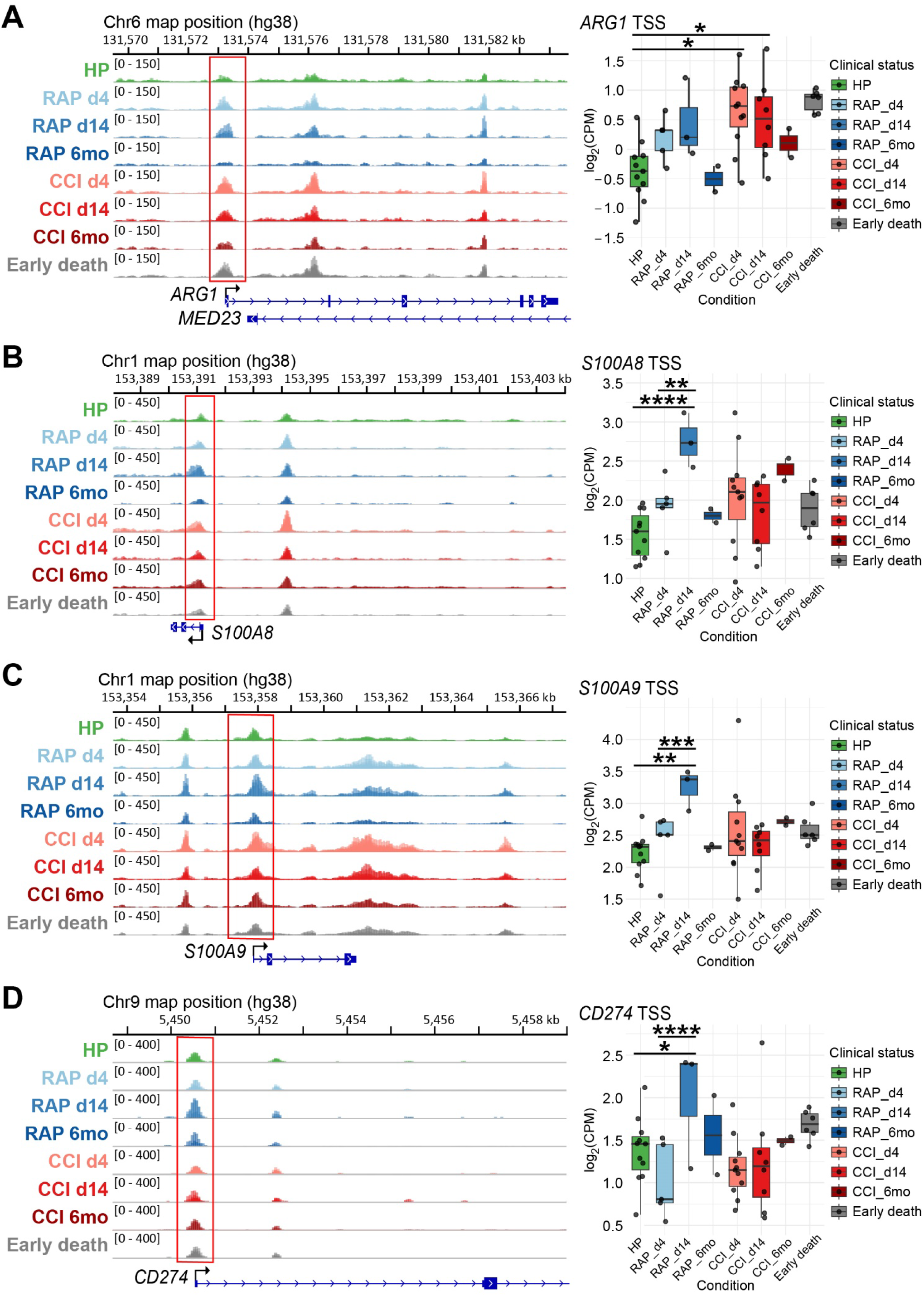
Dynamic changes in chromatin architecture at myeloid-suppressive gene promoters. Genome browser tracks of chromatin accessibility (left) and quantified promoter accessibility (right) in CD66b⁺ MDSCs for (**A**) *ARG1*, (**B**) *S100A8*, (**C**) *S100A9*, and (**D**) *CD274*. Box plots depict log₂(CPM) at promoter-associated ATAC-seq peaks (enclosed in red rectangles). Boxed areas indicate interquartile range, horizontal line indicates the median, and whiskers represent minimum and maximum values (excluding outliers, if any). Samples are grouped and colored by clinical status and time point: green, healthy participant (HP); light, medium, and dark blue, rapid recovery (RAP) d4, d14, and 6mo, respectively; light, medium, and dark red, critical illness (CCI) d4, d14, and 6mo, respectively. Early death patients are colored gray. *, *P* < 0.05; **, *P* < 0.01; ***, *P* < 0.001; ****, *P* < 0.0001.

In contrast, MDSCs from CCI and early-death patients showed little evidence of coordinated, time-dependent chromatin remodeling at these loci. Accessibility at *ARG1* (Fig. 5A) and *S100A8/9* (Fig. 5B, C) remained relatively stable and modestly elevated relative to both HP and RAP at 6 months (Supplementary Table 5), consistent with a constitutively permissive chromatin state. For *CD274*, promoter chromatin remained open across all CCI time points and in early-death samples, without evidence of inducible increases following sepsis onset (Fig. 5D). This stable, non-induced *CD274* chromatin state in CCI, suggests that PD-L1 expression does not undergo stimulus-responsive upregulation. Collectively, these data suggest that immunosuppressive effector genes in CCI reside in a relatively fixed chromatin configuration rather than exhibiting stimulus-responsive regulation.

Other genes frequently upregulated in cancer-associated MDSCs, including *MMP8* and *MMP9* (encoding matrix metallopeptidases), and *NOS2* (encoding inducible nitric oxide synthase), showed no differential promoter accessibility in our dataset (Supplementary Table 5). This indicates that not all canonical MDSC-associated genes are epigenetically engaged in this context.

Consistent with a shift toward an immunosuppressive phenotype, chromatin accessibility at key MHC class II antigen presentation genes was reduced in CD66b⁺ MDSCs from CCI and early-death patients (Fig. 6; Supplementary Table 5). Genome browser tracks revealed significantly decreased accessibility at the *HLA-DRA* promoter (Fig. 6A), along reduced accessibility at *HLA-DRB1* (Fig. 6B). Notably, *HLA-DRB1* also displayed a secondary peak of accessibility near coding exon 2 in HP that was absent in all sepsis samples. Accessibility at the promoter of *CD74* (Fig. 6C), which encodes a critical MHC class II assembly chaperone, was likewise reduced in CCI and early-death patients relative to RAP at d14. This pattern extended across additional MHC class II paralogs, including *HLA-DQA1*, *HLA-DQB1*, *HLA-DPA1*, and *HLA-DPB1* (Supplementary Fig. 6), indicating coordinated repression of the pathway. In contrast to the dynamic increases in accessibility observed at myeloid-suppressive genes in RAP (Fig. 5), these data demonstrate coordinated epigenetic repression of the MHC class II antigen presentation pathway in CCI.

**Figure 6.**
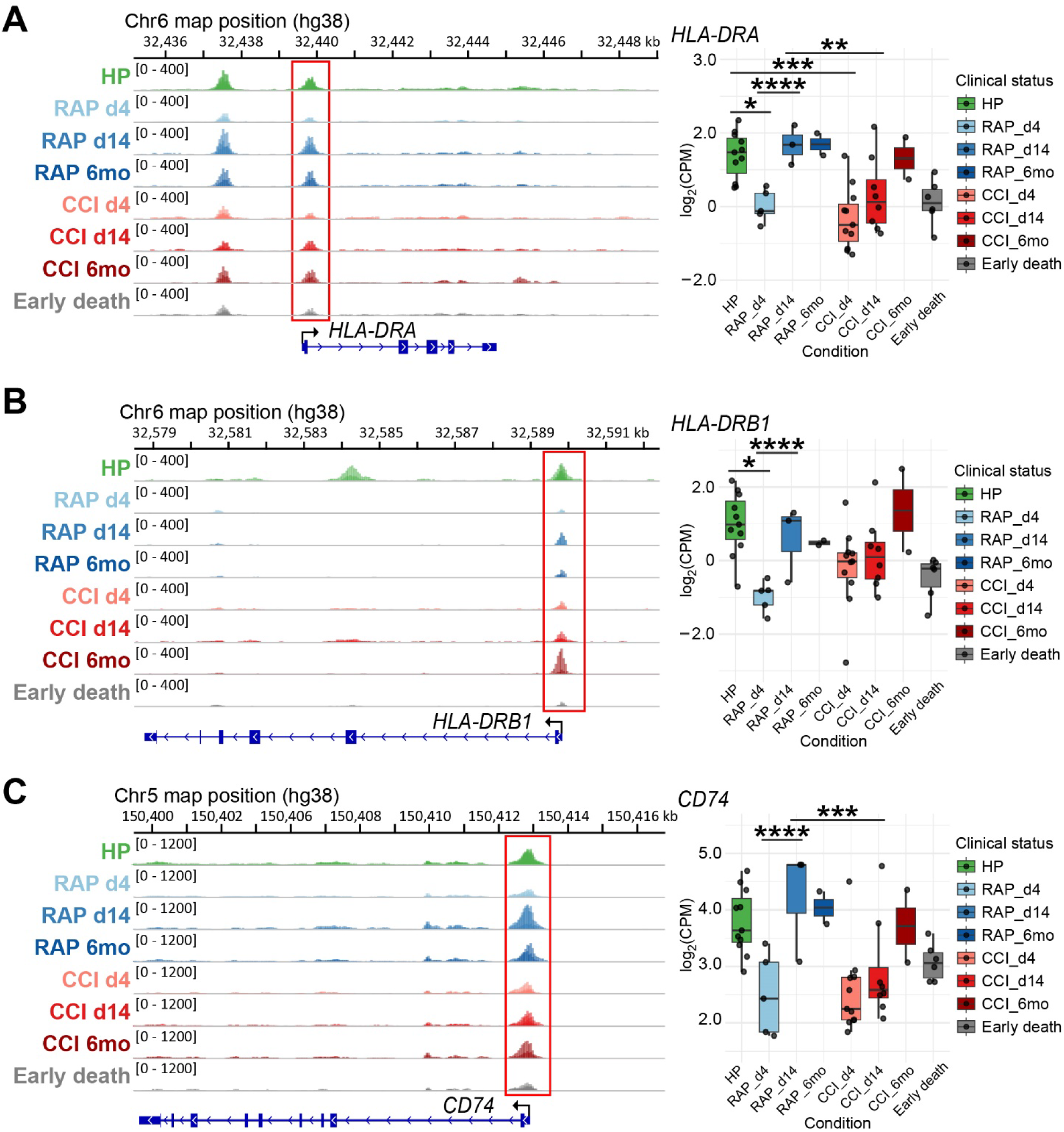
Coordinated epigenetic repression of the MHC class II antigen presentation pathway in chronic critical illness. Genome browser tracks of chromatin accessibility (left) and quantified promoter accessibility (right) in CD66b⁺ MDSCs for representative MHC class II genes, (**A**) *HLA-DRA*, (**B**) *HLA-DRB1*, and (**C**) *CD74*, from HP as well as RAP, CCI, and early death groups. Adjacent HLA loci display background levels of transposase cutting and lack discrete accessibility peaks, supporting specificity of the observed promoter signals. Box plots depict log₂(CPM) at promoter-associated ATAC-seq peaks (outlined in red). Sample group colorations, quantitative box plot values, and other symbols are as in Fig. 5. *, *P* < 0.05; **, *P* < 0.01; ***, *P* < 0.001; ****, *P* < 0.0001.

Whereas MHC class II loci exhibited pathway-level repression in CCI, promoter accessibility across MHC class I antigen processing and presentation genes remained robust in all clinical groups (Supplementary Fig. 7). High levels of accessibility were maintained at loci encoding peptide transporters (*TAP1*, *TAP2*), immunoproteasome components (*PSMB8*, *PSMB9*), the invariant light chain (*B2M*), and the class I heavy chain (*HLA-A*), as well as at *CANX*, which facilitates MHC class I folding. Despite modest fluctuations, these promoters remained strongly accessible across all conditions, indicating preservation of the antigen processing and presentation machinery. Thus, in contrast to the selective repression of MHC class II genes, the MHC class I pathway is epigenetically preserved in CCI.

Further supporting the specificity of these remodeling events, chromatin accessibility at loci unrelated to immune function, including regions within *TMEM72* and upstream of *OTOS* and *FBRSL1*, remained stable across all clinical groups and time points (Supplementary Fig. 8), with no significant changes detected (Supplementary Table 5).

Collectively, these findings indicate that promoter architectures of key immunomodulatory genes in MDSCs are dynamically remodeled in RAP, whereas in CCI and early-death patients, chromatin states are largely fixed, either remaining constitutively open or repressed. This loss of stimulus-responsive chromatin plasticity is coupled by selective repression of the MHC class II pathway alongside preservation of MHC class I antigen processing, defining a rewired and persistently immunosuppressive epigenetic program in CCI.

## Discussion

Building on our previous transcriptomic findings in post-sepsis myeloid compartments (Barrios *et al*., 2024), we hypothesized that persistent MDSC dysfunction in sepsis survivors who do not rapidly recover but instead develop CCI is driven by sustained epigenetic remodeling. Specifically, we posited that CD66b⁺ MDSCs isolated from sepsis survivors would exhibit divergent epigenetic signatures based on time point and clinical outcome (e.g., CCI vs RAP). We further predicted that such reprogramming would be evident in chromatin accessibility profiles at immune-regulatory loci and could be consistent with MDSCs exhibiting distinct functional phenotypes. Our current study tests this hypothesis using high-resolution chromatin profiling of human PBMC CD66b⁺ MDSCs across time and clinical cohorts.

Promoter accessibility at canonical immunoregulatory cytokine loci was limited and largely static across clinical conditions. The *IL10* promoter exhibited a modest but significant increase in accessibility, specifically between RAP d4 and d14, whereas no significant changes were observed at other time points or in CCI patients. In contrast, promoters for *IL10RA* and *IL4R* showed stable accessibility across all cohorts, with *IL10RA* displaying a strong constitutively accessible peak at the TSS and *IL4R* showing only modest baseline accessibility. The *IL4* promoter remained at background levels in all samples, consistent with epigenetic silencing. These findings suggest that classical cytokine-mediated suppressive pathways are not broadly activated through promoter opening in post-septic CD66b⁺ MDSCs. By contrast, promoters of alarmin-associated genes, such as *S100A8* and *S100A9*, remained broadly accessible and increased in accessibility following sepsis, particularly in RAP patients, consistent with sustained activation or reinforcement of inflammatory and myeloid activation programs.

This pattern also may reflect selective reinforcement of pro-inflammatory and tissue-trafficking programs that could exacerbate tissue damage (e.g., with the potential to drive pulmonary injury) in the absence of coordinated adaptive immune recovery. For example, CCI patients at d14 post-sepsis exhibited signatures of sustained inflammation coupled with immune dysfunction, including elevated IL1B-driven prostaglandin synthesis and anti-apoptotic signaling (*via* TNFAIP3/A20) alongside altered MHC-II complex-associated interactions (CD74, HLA-DRA), consistent with impaired antigen presentation. This pattern is indicative of an environment of persistent low-grade inflammation combined with blunted adaptive immunity, a hallmark of sepsis-induced immunoparalysis. Indeed, immunosuppressive myeloid cells (e.g., HLA-DR^lo^ monocytic MDSCs) have been reported to appear early in CCI patients and, by two weeks, exhibit increased inflammatory output (IL1B, etc.) but diminished antigen-presentation capacity, as indicated by downregulated HLA-DRA/HLA-DRB1.

Importantly, our findings refine the concept of sepsis-induced immunoparalysis by demonstrating a striking pathway-level dissociation between MHC class II and MHC class I regulation in CCI. Whereas promoters associated with MHC class II antigen presentation are coordinately repressed, a broad network of genes governing MHC class I antigen processing and presentation, including peptide transporters, immunoproteasome components, and assembly factor, remains strongly accessible. This indicates that impaired antigen presentation in CCI is not due to global chromatin collapse, but instead reflects targeted epigenetic silencing of the MHC class II pathway. In this context, immunoparalysis appears to arise from a failure of adaptive immune priming capacity rather than a general defect in antigen processing. Such selective repression, coupled with preservation of cell-intrinsic antigen processing machinery, provides a mechanistic framework for the coexistence of impaired adaptive immunity and persistent inflammation observed in CCI.

Furthermore, canonical pathway analysis of genes associated with differentially accessible promoter regions between CCI and RAP on day 14 revealed a striking overall suppression of pathway activity in the CCI group. Epigenetic repression of immune, metabolic, and inflammatory programs was evident from numerous signaling pathways with negative *Z* scores (Fig. 4). For example, pathways involved in cytokine signaling (e.g., IL-1, IL-2/IL-7 axes), innate receptor activation (TLR3 signaling), and neutrophil functionality (fMLP signaling, degranulation) were all predicted to be inactivated. This extensive downregulation of critical immune pathways likely underlies the persistence of immature, dysfunctional MDSCs that do not appear to fully transition back to effective antigen-presenting or pro-resolving phenotypes. By contrast, in RAP survivors, these pathways were relatively re-engaged or restored, correlating with the resolution of immunosuppression and a return toward normal myeloid function.

The MDSC epigenome from sepsis survivors with CCI appears “locked” in a state that blunts antigen presentation (e.g., through reduced expression of MHC and co-stimulatory machinery), maintains persistent immunosuppression (e.g., through sustained expression of inhibitory mediators and suppressive receptors), and propagates chronic inflammation via alarmins and oxidative stress. Particularly notable is the strong predicted inhibition of the chromatin modifying enzymes pathway in MDSCs isolated from CCI patients, suggesting a self-reinforcing epigenetic state in which the machinery responsible for remodeling chromatin may itself be repressed. This could limit the capacity of CCI myeloid cells to reset their transcriptional programs, contributing to a persistent immunosuppressive and metabolic phenotype. By contrast, restoration of this activity in RAP patients may be critical for the large-scale promoter opening and immune re-engagement observed during recovery.

Such differences support the possibility that epigenetic reprogramming after sepsis can drive divergent fates, either promoting trained innate immunity or chronic tolerance (Lajqi *et al*, 2023; Wang *et al*, 2024). We speculate that RAP survivors induce beneficial innate immune training that restores immune vigilance and resilience, whereas CCI patients exhibit maladaptive training (mTRIM) or tolerized state that fails to fully reactivate host defenses. Notably, our day 4 data are consistent with an early divergence: patients destined for CCI already show enrichment of pathways dampening lymphocyte activation (implying early innate immune tolerance or exhaustion), whereas RAP patients maintain or even elevate antigen-presentation signals (e.g., higher CD74/HLA-DRA), suggesting preserved innate/adaptive crosstalk. This divergence may contribute to ongoing tissue injury and failed immune resolution in CCI.

Several limitations must be acknowledged relative to these findings. First, our analysis focused exclusively on CD66b⁺ MDSCs from peripheral blood, which may not fully represent tissue-resident myeloid populations that could exhibit distinct epigenetic signatures. Second, our bulk ATAC-seq data do not resolve individual MDSC subtypes, i.e., polymorphonuclear, monocytic, or granulocytic, that may be the primary driver of specific septic immunoregulatory phenotypes. Nonetheless, bulk chromatin accessibility profiling captures population-level regulatory states that are relevant to observed phenotypic MDSC immunosuppression. Third, the relatively small sample sizes, particularly at later time points, limit statistical power for detecting subtle but potentially meaningful chromatin accessibility changes. This includes not having any female sepsis patients classified as RAP at this point in the study, precluding assessment of the contribution of sex. Fourth, chromatin accessibility does not always correlate directly with gene expression or cellular function, and while bulk profiling captures population-level regulatory states, this study did not perform comprehensive paired transcriptomic or functional validation at the single-sample level. Additionally, our study design does not definitively establish causality between observed epigenetic changes and clinical outcomes, as both may reflect downstream consequences of unmeasured variables such as comorbidities, therapeutic interventions, or genetic predisposition. Lastly, the binary classification of patients into CCI versus RAP groups, while clinically relevant, may oversimplify the spectrum of post-sepsis recovery states.

Translation of these findings to clinical practice will require several advances. First, validation in larger, multi-center cohorts will be essential to confirm the reproducibility and generalizability of identified epigenetic signatures across diverse patient populations and healthcare settings. Development of rapid, cost-effective assays to detect key chromatin accessibility markers in clinical laboratories will be necessary for practical implementation. Prospective studies will be required to determine whether early epigenetic profiling (within 4–7 days post-sepsis) can reliably stratify patients at risk for CCI with sufficient sensitivity and specificity to guide clinical decision making. Most importantly, mechanistic studies will be needed to define how epigenetic regulatory pathways are disrupted in CCI and contribute to persistent MDSC dysfunction. These insights will be necessary to inform any future therapeutic strategies aimed at restoring immune competence in post-sepsis patients, which would require extensive preclinical validation and cautious clinical translation. Such efforts should prioritize safety given the immunocompromised state of target patients, while establishing appropriate timing, dosing, and patient selection criteria.

In sum, epigenetic disparities between sepsis survivors who either develop CCI or rapidly recover point to immune training versus immune suppression as a central theme. CCI patients exhibit a stably reprogrammed epigenetic state characterized by selective repression of antigen presentation pathways alongside preservation of core antigen processing machinery. In contrast, RAP patients undergo dynamic immune reprogramming that may facilitate pathogen clearance and tissue recovery. This divergence highlights a loss of chromatin plasticity as a defining feature of poor clinical outcome. Uncovering the mechanism of this bifurcation advances our understanding of sepsis outcomes and therapeutic implications, suggesting that therapeutically skewing the balance towards trained immunity (enhancing monocyte function and antigen presentation) might avert CCI. Identifying these cohort-specific epigenetic states could guide development of precision immunotherapies, including targeted epigenetic therapeutics to rewire tolerized MDSCs and restore immune competence in patients with CCI or prevent its potential onset.

## Methods

### Study design

Our study design was previously reported by Darden et al. (Darden *et al*, 2021a) https://pmc.ncbi.nlm.nih.gov/articles/PMC11076668/. To summarize, this prospective, observational cohort study was registered with *clinicaltrials.gov* (<u>NCT</u>05110937) and conducted at a tertiary care, academic research hospital.

Sepsis was screened using the electronic medical record based Modified Early Warning Signs-Sepsis Recognition System (MEWS-SRS) (Castello & Gavelli, 2024) and confirmed by standard clinical criteria. All patients with sepsis were treated according to sepsis-3 clinical guidelines with early goal-directed fluid administration, broad-spectrum antibiotics, and vasopressor administration if indicated (Castello & Gavelli, 2024; Singer *et al*., 2016). Inclusion and exclusion criteria were:

#### 1. Critically ill sepsis patients

Inclusion criteria:

a. Age ≥18 years
b. Meets criteria for sepsis or septic shock by Sepsis-3 consensus criteria.

Exclusion criteria:

a. Have disease states that predispose to significant immune system dysfunction

i. Have comorbidity burden or goals of care that preclude recovery after sepsis, including
ii. death within 12 hr)
iii. uncontrollable surgical source of sepsis
iv. patients deemed to be futile care or have advanced directives limiting resuscitative efforts
v. alternative diagnoses causing shock state (e.g., hemorrhage, myocardial infarction or pulmonary embolus)
vi. known HIV infection with CD4^+^ count <200 cells/mm^3^
vii. severe traumatic brain injury with unencumbered assessment of Glasgow Coma Scale score equaling 3 on admission to the intensive care unit.
b. known pregnancy
c. enrollment >96 hr after suspected sepsis onset
d. pre-hospitalization bedridden performance status (WHO/Zubrod score ≥4)

i. subsequent clinical adjudication diagnosis not consistent with sepsis/septic shock by Sepsis-3 criteria.
e. burn Injury greater than 20% Total Body Surface Area

#### 2. Healthy participants (control population)

Inclusion criteria:

a. All adults (age ≥18)
b. Ability to obtain Informed Consent prior to blood collection.

Exclusion criteria:

a. Current, chronic steroid use
b. Known pregnancy
c. Current or recent (within 7 days) use of antibiotics.

CCI was defined as ICU length of stay ≥14 days with persistent organ dysfunction as measured by the Sequential Organ Failure Assessment (SOFA) score. Patients were also designated CCI with <14 days ICU length of stay if they were transferred to another hospital, or discharged to a long-term acute care facility or hospice with evidence of persistent organ dysfunction (Castello & Gavelli, 2024; Hollen *et al*, 2019; Mathias *et al*, 2017b).

Whole blood samples were collected from 19 patients at 4 days (5 RAP, 11 CCI, 3 early death), 14 at 2-3 weeks (3 RAP, 8 CCI, 3 early death), and 4 at 6 months (2 RAP, 2 CCI) after initiation of sepsis protocols (Supplementary Table 6). Additionally, 11 samples were obtained from healthy participants (Supplementary Table 3). Differences in age and sex distributions among HPs, RAP, and CCI groups at each time point were assessed using ANOVA and chi-square tests, respectively (Supplementary Table 6).

### Cell isolation

Whole blood from ethylenediaminetetraacetic acid-coated blood collection tubes was diluted 1:1 with phosphate-buffered saline (PBS) and layered over Ficoll Paque PLUS (GE Healthcare, Chicago, IL) to isolate PBMCs through centrifugation. PBMCs were resuspended in PBS with 5% (v/v) fetal calf serum before isolating CD66b^+^ cells with “The Big Easy” EasySep™ magnet (STEMCELL Technologies, Vancouver, BC, Canada) using EasySep HLA Whole Blood CD66b Positive Selection Kit (STEMCELL Technologies, #17882). Isolated cells were stained with anti-CD66b conjugated to BV421 and analyzed by flow cytometry. Isolation kits return >80% purity of low- and high-expressing CD66b^+^ cells (Barrios *et al*., 2024; Darden *et al*., 2021a).

Total CD3^+^ lymphocytes were isolated with magnet using EasySep™ Human T Cell Isolation Kits (STEMCELL Technologies, #17951), according to the manufacturer’s instructions.

### T-cell suppression assay

Suppression of activated T-lymphocyte proliferation was performed as described previously (Barrios *et al*., 2024). Briefly, isolated CD3^+^ T cells were labeled with cell trace violet (Thermo Fisher, Waltham, MA, #C34557), and 1 x 10^5^ cells were plated without (unstimulated, Unstim control) and with anti-CD3/CD28 antibodies (Stim) (STEMCELL Technologies, #10971). To measure T-cell proliferation suppression, an equal number of CD66b^+^ cells were added for co-culture for 4 days at 37°C and 5% CO_2_. Then, cells were harvested, stained with anti-CD8 FITC and anti-CD4 PE, and analyzed via flow cytometry (ZE5 Cell Analyzer, Bio-Rad Laboratories, CA) to calculate proliferation indices. To test for significant differences between sample stratified by clinical status, we fit two-way ANOVA tests with patient groups (RAP vs CCI) and stimulation conditions. Tukey’s post-hoc pairwise comparisons were performed using the estimated marginal means.

### Omni-ATAC and library preparation and sequencing

Aliquots of 100,000 CD66b^+^ MDSCs were centrifuged at 500 x *g* for 5 min at 4°C. Cell pellets were resuspended in 150 µL Bambanker Freezing Media (Bulldog Bio cat# BB05) and stored at −80°C until use. Frozen cell aliquots were thawed on ice by adding 800 µl of cold wash buffer (10 mM Tris-HCl, pH 7.4, 10 mM NaCl, 3 mM MgCl_2_, 0.1% (w/v) Tween-20) (Corces *et al*., 2017). Cells were centrifuged at 500 x *g* for 5 min at 4°C and resuspended into 500 µL of cold wash buffer for cell counting. Omni-ATAC reactions were performed with 75,000 cells, spiked with 8,000 *Drosophila* ATAC-Seq Spike-In Control nuclei (Active motif cat# 53154). Human cells and *Drosophila* spike-in mixture were centrifuged at 500 x *g* for 5 min at 4°C, and the pellets were resuspended into 50 µL cold lysis buffer (wash buffer with 0.1% (w/v) NP40 and 0.05% (w/v) digitonin) (Corces *et al*., 2017). The cell and nuclei mixes were lysed on ice for 6 min, then lysis reaction was stopped by adding 1 mL of cold wash buffer. The nuclei were centrifuged at 1,000 x *g* for 10 min at 4°C and the pellets were resuspended in 50 µL Tagmentation buffer (Zymo-Seq ATAC Library Kit cat# D5458), supplemented with 0.05% (w/v) digitonin and 0.1% Tween-20. We found that this formulation consistently outperformed other sources.

Tagmentation reactions and library preparation were conducted according to the manufacturer’s instructions for the Zymo-Seq ATAC Library Kit. Final libraries were double-size-selected using AMPure beads to enrich for fragments between 150 bp and 500 bp. Final Omni-ATAC libraries were submitted to the UF Interdisciplinary Center for Biotechnology Research (UF-ICBR, RRID:SCR_019152) NextGen Sequencing Core for sequencing on an Illumina Novaseq X 10B at 2 x 150 nt targeting between 100 and 150 million reads per library.

### Omni-ATAC data processing

FASTQ files from 48 samples were processed using the nf-core (Ewels *et al*, 2020) atacseq pipeline (v2.1.2) in Nextflow (v22.10.1) (Di Tommaso *et al*, 2017) with default parameters unless specified otherwise. Briefly, raw FASTQ reads were quality checked with FastQC (v0.11.9) (Andrews, 2010) and adaptor trimmed using Trim-Galore (https://www.bioinformatics.babraham.ac.uk/projects/trim_galore/) (Martin, 2011). Reads were aligned to a combined reference genome of human (GRCh38, NCBI refSeq annotation gtf) and *D. melanogaster* (Dmel) (BDGP6, NCBI refSeq annotation gtf) using BWA-MEM (v0.7.17) (Li & Durbin, 2009) with paired-end settings (-M flag). BAM files were filtered for properly paired reads (samtools v1.12, -f 2), and duplicates were removed using Picard MarkDuplicates (v2.25.0). Mitochondrial reads were excluded to focus on nuclear chromatin.

For browser track visualization, bigwig files were generated from human mapped reads using deepTools (v3.5.1) (Ramírez *et al*, 2014) bamCoverage. Normalization of counts was performed by utilizing a scaling factor calculated using the median Dmel count values. Consensus peaks were called across biological replicates using MACS2 (v2.2.7.1) (Zhang *et al*, 2008) (Gaspar, 2018) within the pipeline, with a *Q* value threshold of 0.05 and Tn5 insertion offset correction.

Differential accessibility analysis was performed using the consensus peak feature counts matrix for human peaks in R (v4.5.1), using edgeR (v4.6.3) (Robinson *et al*, 2010), limma (v3.64.1) (Ritchie *et al*, 2015), and voom(Law *et al*, 2014). The 48 Omni-ATAC samples in this study were processed across 7 batches. To correct for this, a scaling factor was derived from the median Dmel counts and used to adjust the library size for each sample (norm.factors were then set to 1). For the design matrix, a new variable was created, condition, consisting of the samples time point, e.g., d4, d14, and the clinical status, e.g., RAP. The design matrix included batch and condition for male only samples, or batch, sex, and condition for both sexes as covariates. Peaks with low accessibility (CPM < 1 in 70% of the smallest group (condition)) were filtered out.

The voom method was used to model mean-variance relationships by generating precision weights for each observation. Sample counts were then analyzed with limma by fitting the linear model design = ∼ batch + condition, with patient included as a blocking factor to account for repeated measures. Within-subject correlation was estimated using limma’s duplicateCorrelation function (consensus correlation = 0.24) and incorporated into the voom/lmFit workflow. Differential tests were performed using the empirical Bayes moderated *t*-statistics (eBayes) on the contrasts of interest. This approach applied precision weights to the log_2_ counts per million (log_2_(CPM)) in a weighted least squares framework, testing for differential expression between condition groups while moderating gene-wise variances across all genes. *P* values were adjusted globally (across all contrasts, using the Benjamini-Hochberg method, and peaks with a global FDR < 0.01 and an absolute log_2_ fold change of at least 1.0 (2x fold change) were considered significantly differentially accessible. Differential peaks were annotated using HOMER (Heinz *et al*, 2010).

### PCA and UMAP plot generation

For visualization of high-dimensional Omni-ATAC data, principal component analysis (PCA) was performed using adjusted log_2_(CPM) data, using the removeBatchEffect function, with batch, and sex if needed, as batches, plus condition. The top 1,000 most-variable peaks across samples were used for both. For the two healthy participants with repeated blood draws, the PCA and UMAP results were merged into a single data point for clarity (*n* = 46). The first 30 principal components (PCs) were selected as the input for Uniform Manifold Approximation and Projection (UMAP) to generate two-dimensional embeddings for sample visualization. UMAP and PCA plots were generated in R using custom in-house scripts for ggplot2 based plots. UMAP plots were generated using the R package uwot.

### Gene Ontology enrichment analysis

Gene Ontology (GO) enrichment analysis was performed using the clusterProfiler (Wu *et al*, 2021; Yu *et al*, 2012) R package. Peaks were used for GO enrichment if they met the statistical significance above and were annotated to be within ±1 kb of a TSS. For the background peak list, all peaks within ±1 kb of a TSS that passed filtering for low counts were used, regardless of statistical significance. A hypergeometric test was used to obtain *P* values, which were adjusted for multiple hypotheses testing using the false discovery rate (FDR) method with 5% FDR. Plots were generated in R using a combination of enrichplot (Yu) and in-house scripts.

### Ingenuity Pathway Enrichment Analysis

Ingenuity Pathway Analysis (IPA) of Differentially Accessible Regions (DARs) from the Omni-ATAC analysis of CD66b⁺ MDSCs were identified by comparing the CCI group to the RAP group at d14 post-sepsis. We restricted this analysis to peaks overlapping gene promoter regions (i.e., ±1kb of transcription start sites) to focus on regulatory elements directly controlling gene expression. IPA (Qiagen, USA) was then performed on the gene list derived from these promoter DARs (filtered at global adjusted *P* < 0.01 and an absolute log_2_ fold change of at least 1 for differential accessibility). In IPA, a Core Analysis of Canonical Pathways was run using the genes associated with the significant promoter DARs as input. This analysis provided both pathway enrichment significance (-log *P* values) and activation *Z* scores, predicting pathway activation or inhibition in CCI relative to RAP. All analyses were conducted using default settings for human datasets, allowing identification of enriched pathways and their predicted activation states based on the promoter accessibility differences between CCI and RAP MDSCs.

### Data processing for box plots

For differential accessibility comparisons between individual genes/peaks (Fig. 5), box plots were generated from Dmel scaled counts that were converted to CPM and plotted as log_2_(CPM) on the y-axis. This ensured that the data used for the IGV tracks and boxplots were processed in a similar fashion. The globally adjusted *P* values from the differential accessibility analysis, described in Omni-ATAC data processing methods section, were added to the box plots. For global accessibility pattern analysis (Supplementary Fig. 5), effect size analysis was employed due to the large number of data points, where traditional significance testing can detect statistically but not biologically meaningful differences. Cohen’s *d* was calculated as the standardized mean difference between groups, with values of 0.2, 0.5, and 0.8 representing small, medium, and large effects, respectively. Mean differences in log₂ CPM values were also calculated, where a difference of 1.0 represents a 2-fold change in accessibility.

## Data availability

All data associated with this study are present in the paper or the Supplementary Materials.

## Acknowledgments

The authors thank the following people for their role in patient recruitment and retention, data collection, and sample collection: LaShaun Bryant, BS; Brandi Buscemi, AS – Physical Therapist Assistant; Ruth Davis, BSN; Jennifer Lanz, MSN, RN; Ashley McCray, ASN; and Ivanna Rocha, MPH. This work was supported, in part, by the grants from the National Institutes of Health, RM1 GM139690 (MPK, CEM, PAE, LLM), R35 GM140806 (PAE), NIH T32 GM008721 (CER, MHR, WBW, AMC) and by the University of Florida Health Cancer Institute. The UF Health Cancer Institute is supported in part by state appropriations provided in <u>Fla. Stat.</u> <u>§ 381.915</u> and the NIH National Cancer Institute under Award Number P30 CA247796. The content is solely the responsibility of the authors and does not necessarily represent the official views of the National Institutes of Health or the State of Florida.

## Author contributions

J.O.B. and M.-P.L.G. performed experiments, analyzed data, and contributed to data interpretation. R.W., M.H.-R., L.Z.-S., M.L.D., R.F.U., I.L.R., W.B.W., A.M.C., F.X., L.E.B., A.M.M., S.D.L., J.C.R., S.M.W., M.A.B., and T.J.L. contributed to sample processing, experimental execution, and data acquisition. J.O.B., G.C., C.E.R., and R.B. performed computational analyses and assisted with data interpretation. L.L.M., C.E.M., and P.A.E. contributed to experimental design, data interpretation, and provided critical resources. P.A.E. and M.P.K. conceived and oversaw the study. J.O.B., M.-P.L.G., and C.E.R. drafted the manuscript with input from all authors. All authors reviewed, edited, and approved the final manuscript.

## Disclosure and competing interests statement

The authors declare that they have no conflict of interest.

## Expanded view content

### This PDF includes

Figs. S1 to S8

Tables S1 to S6

---

| **Contrasts** | **# DARs** | **Increased Accessibility** | **Decreased Accessibility** |
| --- | --- | --- | --- |
| **A. CCI_d4_vs_HP** | 104 | 53 | 51 |
| **B. RAP_d4_vs_HP** | 42 | 8 | 34 |
| **C. CCI_d14_vs_HP** | 72 | 65 | 7 |
| **D. RAP_d14_vs_HP** | 3,900 | 3,897 | 3 |
| **E. CCI_d14_vs_CCI_d4** | 1 | 1 | 0 |
| **F. RAP_d14_vs_RAP_d4** | 22,970 | 22,970 | 0 |
| **G. CCI_d4_vs_RAP_d4** | 61 | 58 | 3 |
| **H. CCI_d14_vs_RAP_d14** | 2,905 | 16 | 2,889 |

## Notes

### Competing Interest Statement

The authors have declared no competing interest.

### Summary of Updates

All sections have been edited. In particular, data and discussion of promoters of genes encoding MHC-I and MHC-II products have been added; Supplemental files updated.

