## Supplemental Figures for "Divergent chromatin remodeling in post-sepsis MDSCs underlies MHC class II repression in CCI"

### Supplemental Material

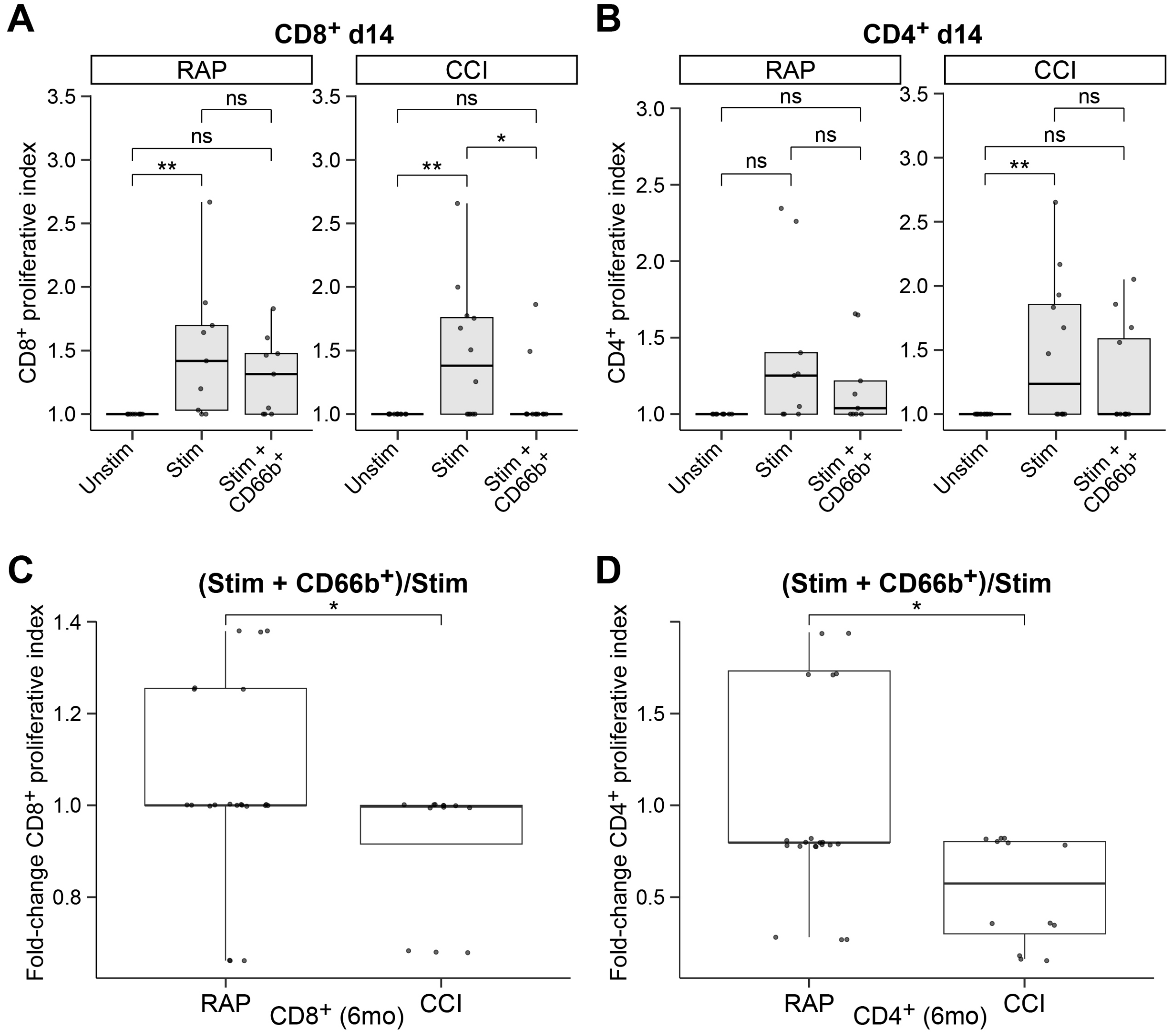


**Figure S1. PMBC CD66b⁺ cells isolated from sepsis survivors demonstrate suppression of T-cell proliferation, confirming MDSC activity.** Total CD3⁺ T lymphocytes were not stimulated (Unstim control) or stimulated with anti-CD3/CD28 antibodies (Stim) and co-cultured with CD66b⁺ cells (Stim + CD66b⁺) isolated from circulating PBMCs obtained from rapid recovery (RAP) or chronic critical illness (CCI) sepsis survivors at the indicated times (d14 or 6mo post-sepsis diagnosis). Box plots of proliferative indices of (**A**) CD8⁺ T cells and (**B**) CD4⁺ T cells relative to Unstim control T cells alone or as fold change of Stim + CD66b⁺ (RAP or CCI) normalized to Stim of (**C**) CD8⁺ T cells and (**D**) CD4⁺ T cells. Box areas indicate the interquartile range (IQR), horizontal lines are the median, and whiskers extend to the most extreme non-outlier values within 1.5 × IQR. Dots represent individual sample proliferative indices in **A** and **B** and fold changes in proliferative indices in **C** and **D.**

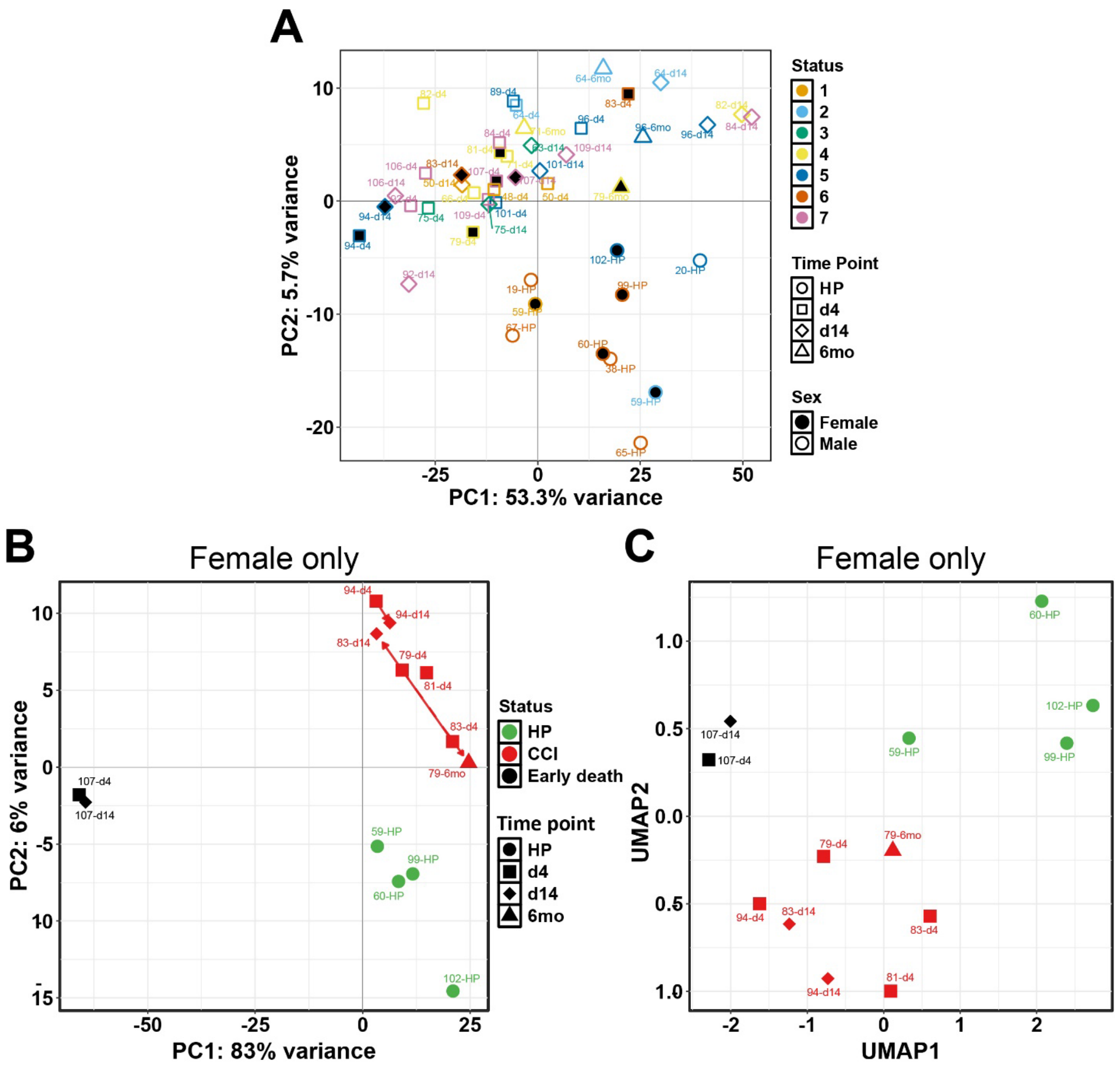


**Figure S2. Dimensionality reduction plots of female-only cohort. A** PCA plot of chromatin accessibility data for all samples (as in Fig. 1A). Points are colored by Omni-ATAC processing batch, shape indicates time point, with filled shapes indicating female, empty shapes for male, participants. **B** PCA plot of chromatin accessibility data (top 1,000 most-variable peaks) from female-only cohort of healthy participants (HP), sepsis survivors who developed chronic critical illness (CCI), and early death (patients who died from septic shock <14 days of entry into intensive care unit). Points are colored by clinical status (HP, CCI, Early death) and shaped by time point. Text labels are sample IDs; arrows show within-patient trajectories: solid arrow = d4 → d14. Arrow direction indicates the temporal progression of each patient in PCA space; arrow length is the Euclidean displacement of the projected samples. Group sizes: HP (n = 4), CCI (n = 7), Early death (n = 2). **C** UMAP plot of the same female cohort as in **B**. The first 30 PCA dimensions for the top 500 most-variable peaks were used for UMAP. Number of nearest neighbors = 4, minimum distance = 0.01. Plot is labeled as in **A** and **B**. Key is as in **B**.

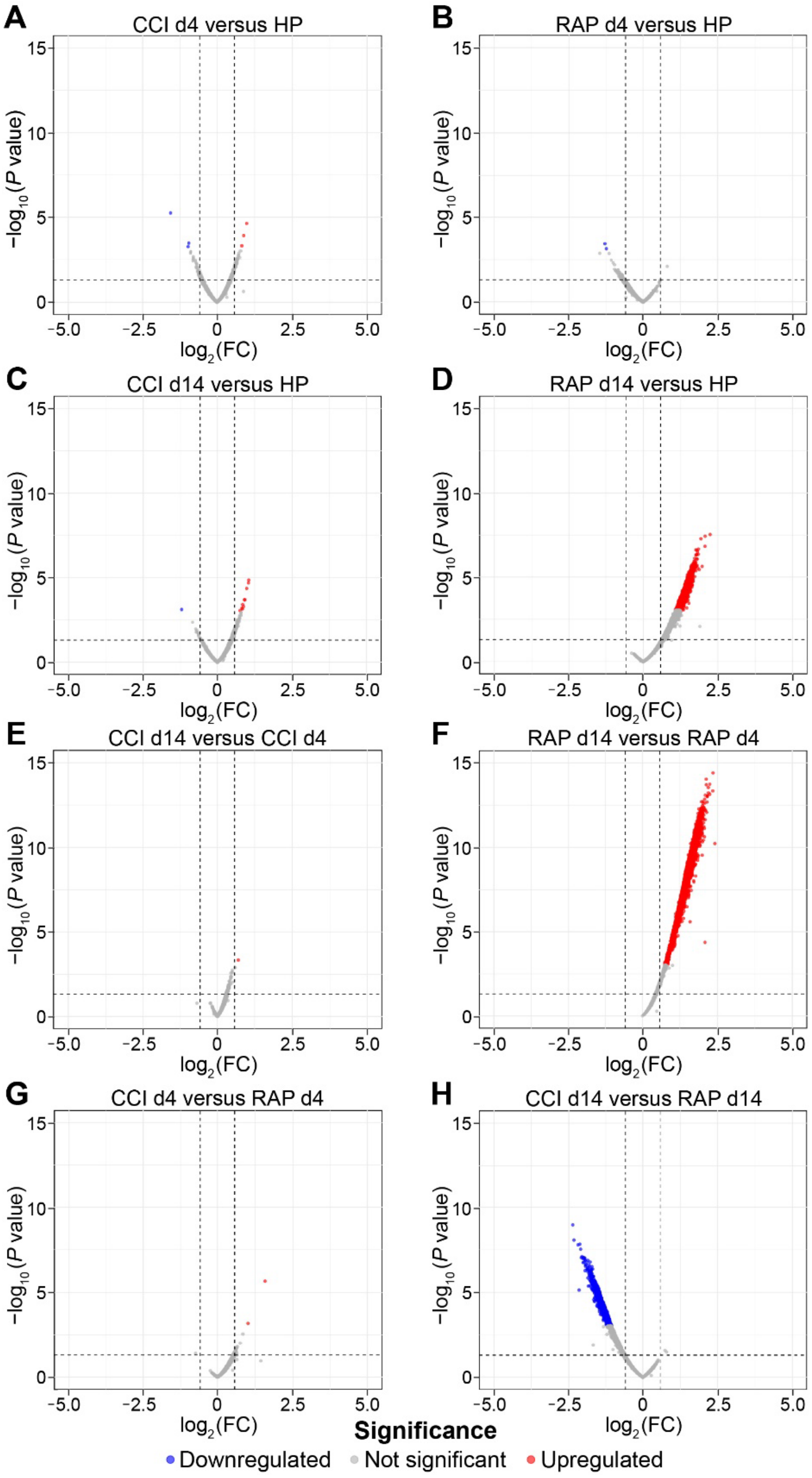


**Figure S3. Volcano plots of differential chromatin accessibility at promoters comparing clinical status and time point within and across groups.** RAP and CCI clinical groups at d4 and d14 versus HP (**A-D**), temporal changes within CCI and RAP across time (**E** and **F**) and between group comparisons for CCI and RAP at d4 and d14 (**G** and **H**). Each DAR is plotted by the -$\log_{10}$ of raw *P* value on y-axis and $\log_2$ Fold Change on the x-axis. DARs were considered significant if the global adjusted *P* value was ≤ 0.01 and $\log_2$ Fold Change was ≥ 1.0. Significant DARs are colored blue for decreased accessibility and red for increased accessibility. Gray indicates no change in accessibility.

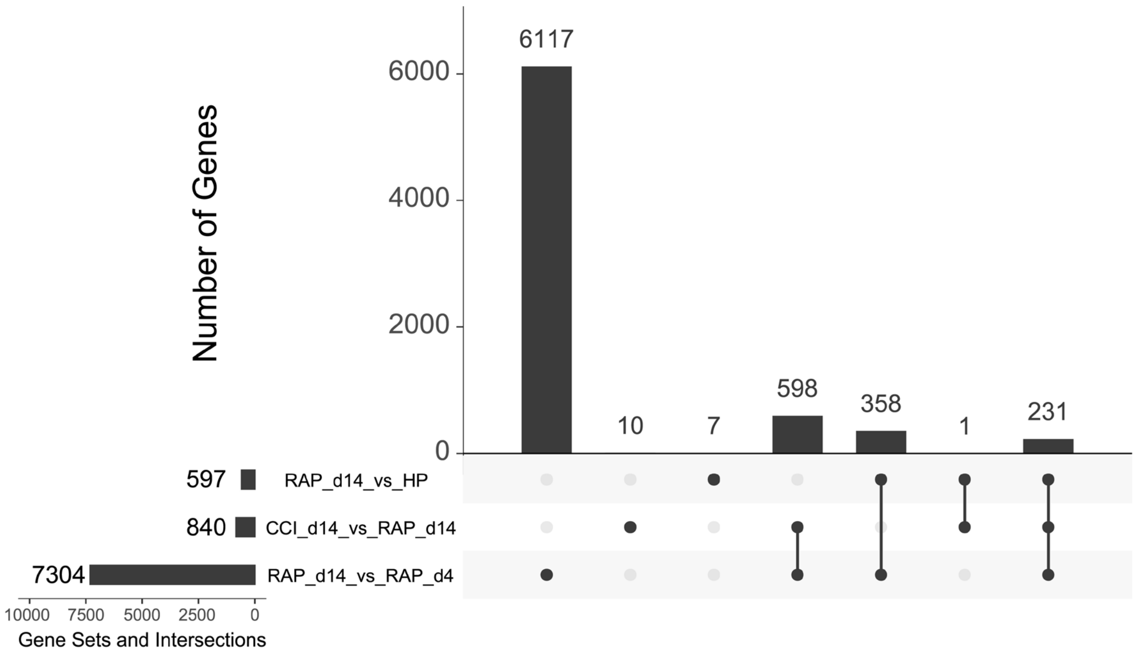


**Figure S4. Unique and common genes associated with promoter DARs between group contrasts.** UpSet plot illustrating the overlap of genes with promoter-associated differentially accessible regions (DARs) across three contrasts: (1) RAP d14 versus HP, (2) CCI versus RAP at d14, and (3) RAP d14 versus RAP d4. The vertical bars represent the number of genes unique to a single contrast (indicated by a single black dot) or shared among multiple contrasts (indicated by connected black dots). The horizontal bar chart shows the total number of genes with promoter DARs in each individual contrast.

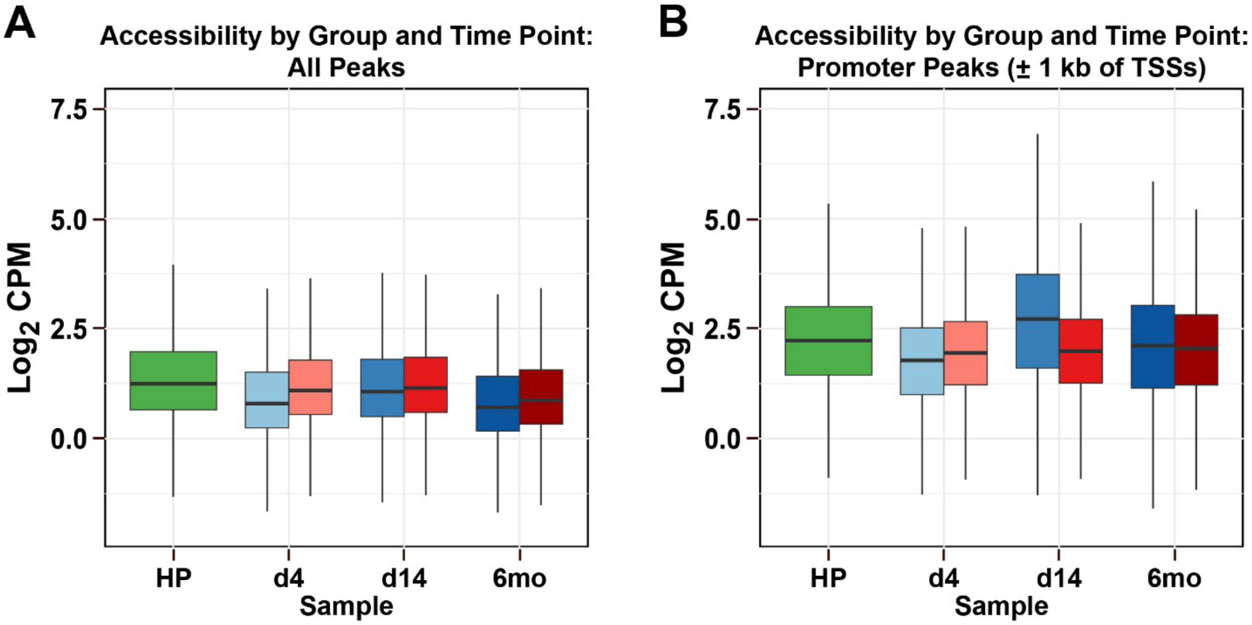


**Figure S5. Dynamic global changes in Omni-ATAC promoter peaks by clinical status across time.** Box plots were generated from $\log_2$ CPM data for all peaks (**A**) and promoter peaks (**B**). The box area indicates the interquartile range, horizontal line is the median, and the whiskers represent minimum and maximum values (excluding outliers, if any). Effect sizes (Cohen's *d*), small (0.2), medium (0.5), large (0.8+) for all contrasts are in Table S2. *P* values not shown due to large sample sizes; focus on biological significance via effect sizes. Samples are grouped and colored by clinical status and time point: green, healthy participant (HP); light, medium, and dark blue, rapid recovery (RAP) d4, d14, and 6mo, respectively; light, medium, and dark red, critical illness (CCI) d4, d14, and 6mo, respectively.

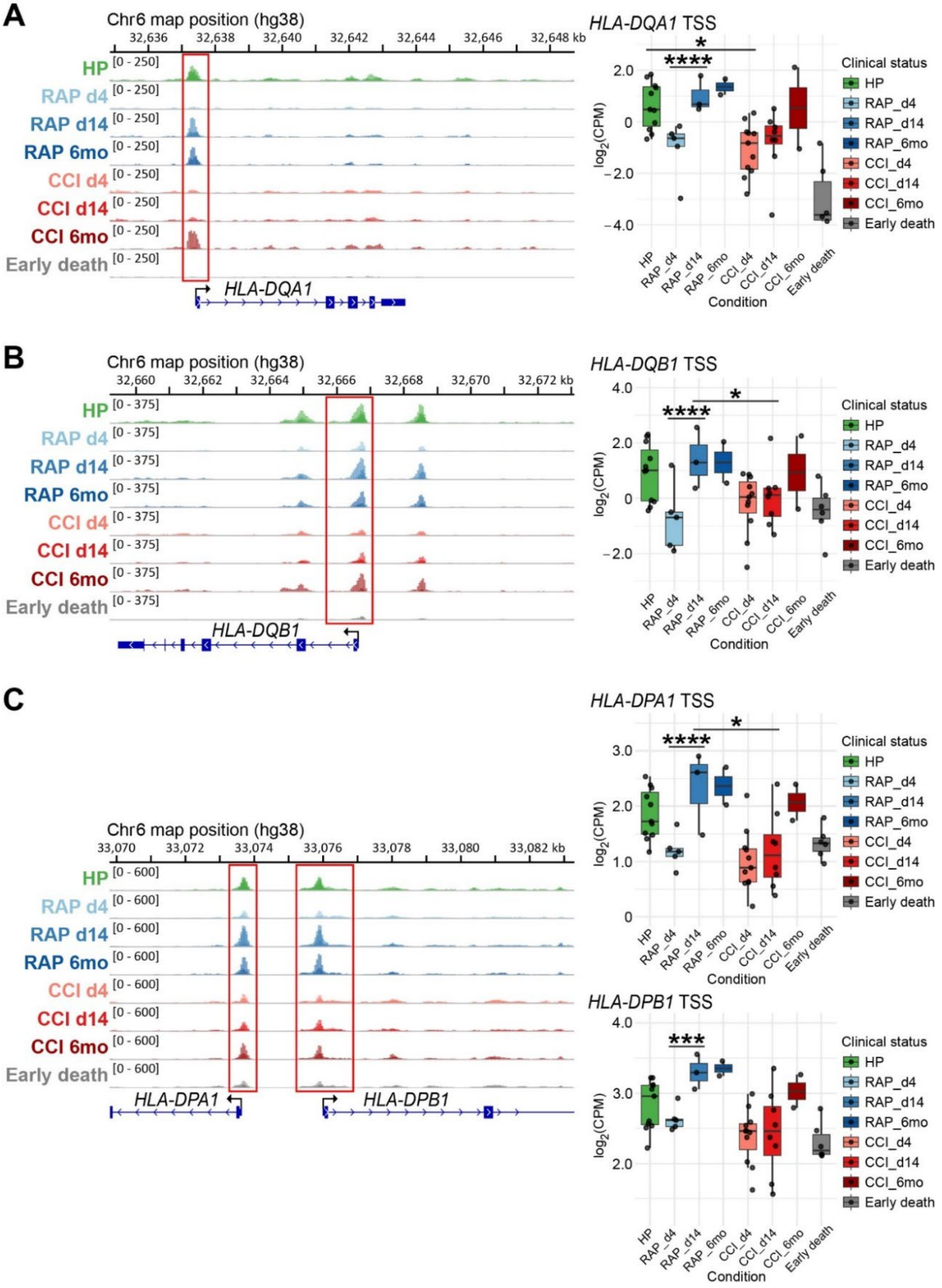


**Figure S6. Consistent repression across MHC class II paralogs reinforces pathway-level epigenetic silencing.** Genome browser tracks of chromatin accessibility (left) and quantified promoter accessibility (right) in CD66b$^+$ MDSCs for (**A**) *HLA-DQA1*, (**B**) *HLA-DQB1*, and (**C**) divergently transcribed *HLA-DPA1*-*HLA-DPB1* locus. Box plots depict $\log_2$(CPM) at promoter-associated ATAC-seq peaks (enclosed in red rectangles). Sample group colorations, quantitative box plot values, and other symbols are as in Fig. 5. *, $P < 0.05$; **, $P < 0.01$; ***, $P < 0.001$; ****, $P < 0.0001$.

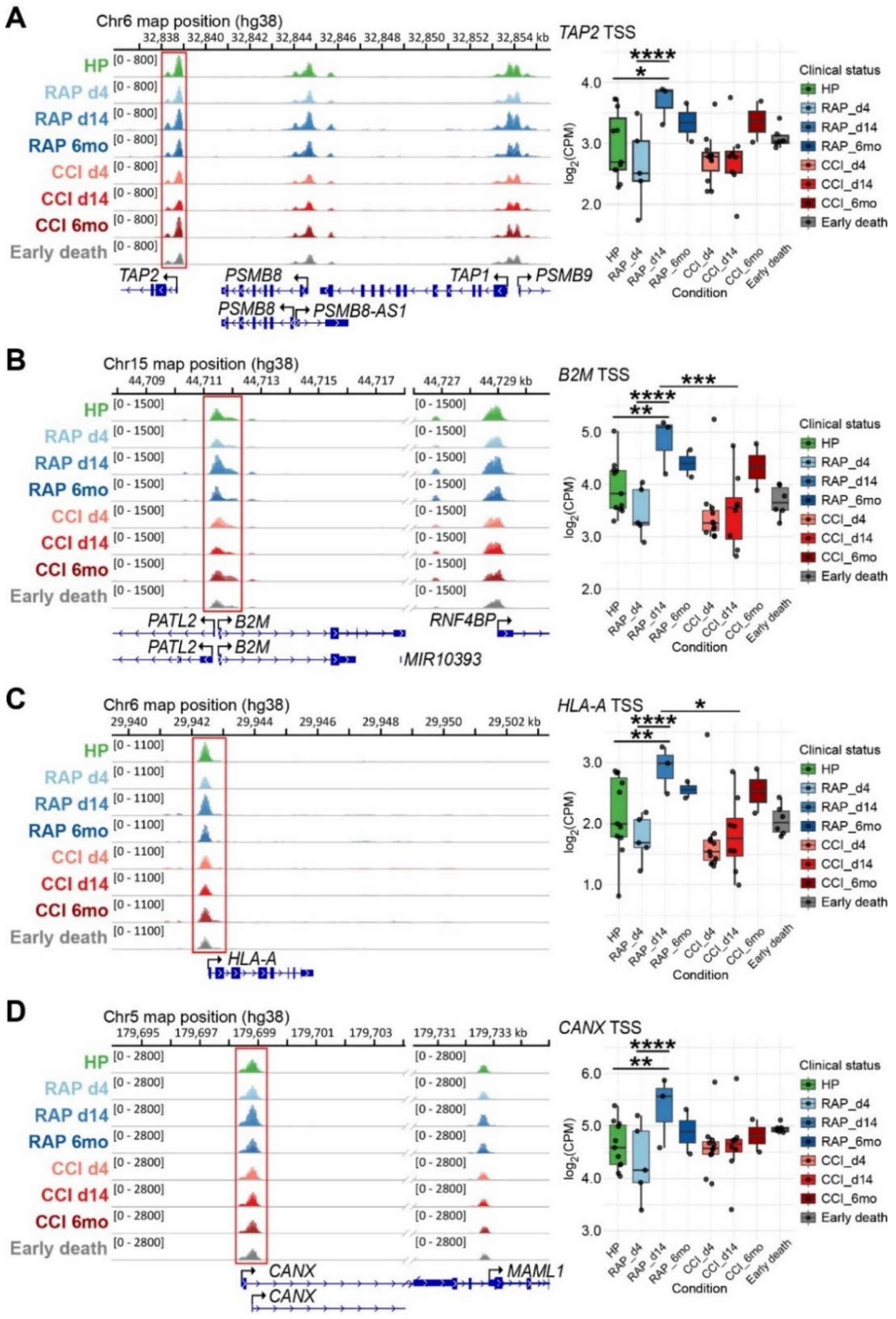


**Figure S7. Preserved antigen processing and MHC class I promoter accessibility contrasts with selective repression of the MHC class II pathway.** Genome browser tracks of chromatin accessibility (left) and quantified promoter accessibility (right) in CD66b$^{+}$ MDSCs for (**A**) the *TAP2–PSMB8–TAP1–PSMB9* locus, with quantification corresponding to the *TAP2* promoter peak, (**B**) *B2M*, (**C**) *HLA-A*, and (**D**) *CANX*. Discontinuous genomic regions (indicated by //) are shown in **B** and **D** to include nearby downstream loci (*RNF4BP* and *MAML1*, respectively), providing internal context for accessibility patterns while preserving consistent scaling across panels. Box plots depict $\log_2$(CPM) at promoter-associated ATAC-seq peaks (enclosed in red rectangles). Sample group colorations, quantitative box plot values, and other symbols are as in Fig. 5. *, $P < 0.05$; **, $P < 0.01$; ***, $P < 0.001$; ****, $P < 0.0001$.

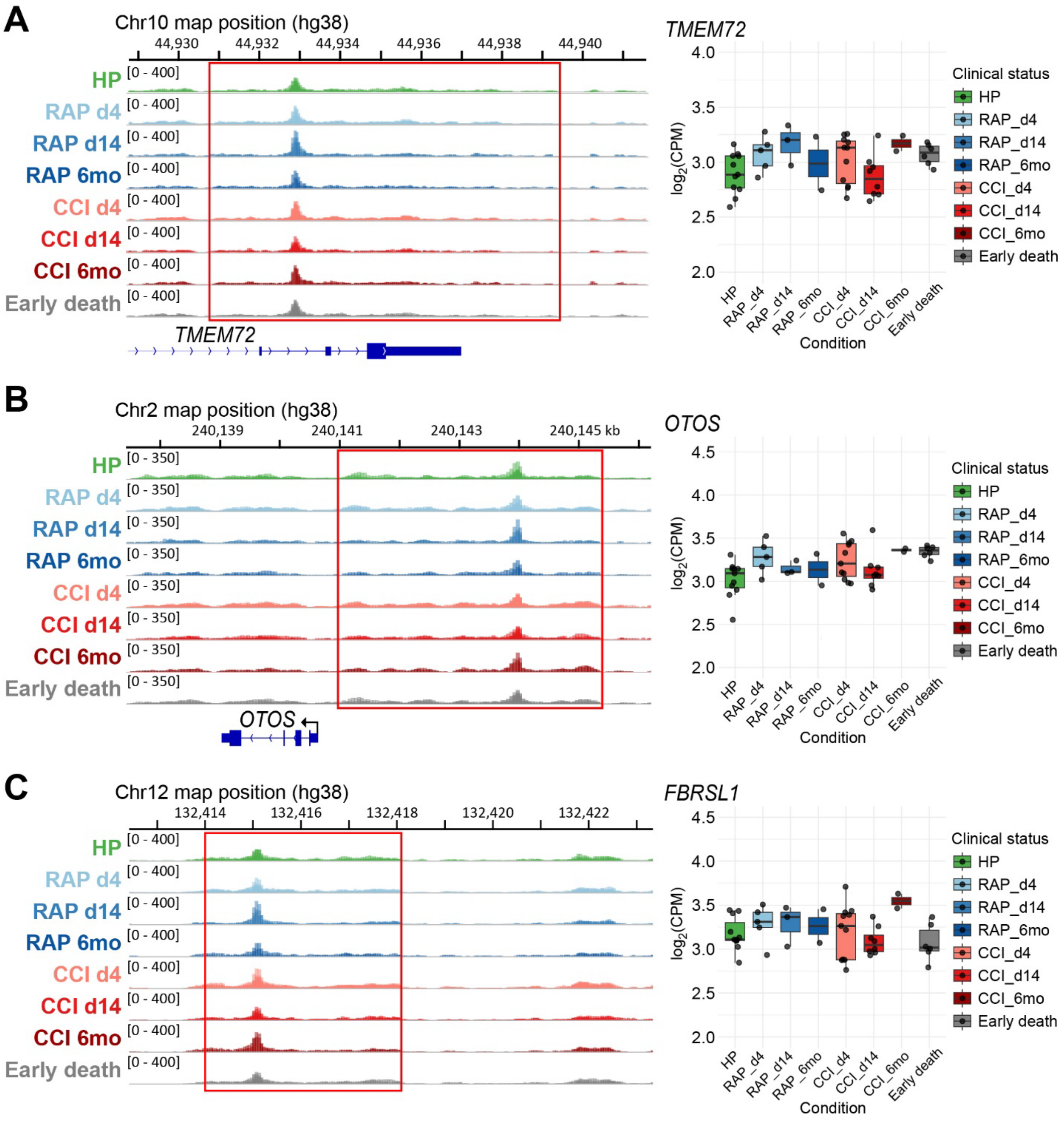


**Figure S8. Stable chromatin accessibility at immune-unrelated loci supports specificity of pathway-selective regulation.** Genome browser tracks of chromatin accessibility (left) and quantified promoter accessibility (right) in $CD66b^+$ MDSCs for (**A**) *TMEM72*, (**B**) *OTOS*, and (**C**) *FBRSL1*. Box plots depict $\log_2$(CPM) at promoter-associated ATAC-seq peaks (enclosed in red rectangles). Sample group colorations, quantitative box plot values, and other symbols are as in Fig. 5. None of the eight comparisons performed in Supplementary Table 5 met significance thresholds.

**Table S1. Differentially accessible regions (DARs) with genomic annotations.** Each row represents a consensus ATAC-seq peak tested for differential chromatin accessibility. Genomic coordinates are reported as chr:start-end (0-based, BED format) in the **Unique_ID** column. Statistical results were derived from limma-voom analysis: **logFC**, $\log_2$ fold-change between conditions; **AveExpr**, average $\log_2$ CPM expression across all samples; **t**, moderated t-statistic; **B**, log-odds that the region is differentially accessible; **P.Value**, nominal p-value; **adj.P.Val**, Benjamini-Hochberg adjusted p-value within contrast; **adj.P.Val.Global**, Benjamini-Hochberg adjusted p-value computed globally across all contrasts. Genomic annotations were generated using HOMER annotatePeaks.pl and include the nearest gene (**Gene.Name**), distance to the nearest transcription start site (**Distance.to.TSS**), and detailed feature annotation (**Detailed.Annotation**). **Simple_Annotation** provides a simplified feature category (e.g., promoter, exon, intron, intergenic) derived from the HOMER detailed annotation.

**Table S2. Comparisons of effect size of dynamic chromatin accessibility changes by clinical status across time between global and promoter peaks.** Table of effect size calculation results for promoter only peaks (Sheet 1; promoter_peaks) and globally for all peaks (Sheet 2; all_peaks). Column headers: **group1**: first condition for contrast; **group2**: second condition for contrast; **Cohens_d**; Cohen's *d* score calculated for contrast defined by group1 and group2; **median_difference**: median of $\log_2$ CPM difference between group1 and group2; **mean_difference**: mean $\log_2$ CPM difference between group1 and group2 contrast: **mean_group1** and **mean_group2**: means of $\log_2$ CPM values for group 1 and group 2, respectively. Cohen's *d* value of 0.2, 0.5, and 0.8 represent small, medium, and large effect sizes, respectively.

**Table S3. Enriched Gene Ontology terms for promoter DARs.** Table of enriched GO terms from promoter DARs. Rows are sorted by FDR adjusted *Q* values. Column definitions: **ONTOLOGY**: GO term ontology category; biological process (BP), cellular component (CC), and molecular function (MF); **ID**: Official GO ID; **Description**: Name of the GO term; **GeneRatio**: Ratio of affected genes in contrast and total genes in the GO term; **BgRatio**: Ratio of the number of genes in experimental dataset background/universe and total number of genes in background/universe; **RichFactor**: Enrichment factor, calculated as GeneRatio/BgRatio; 1 = no enrichment, > 1 = enriched, < 1 depleted; **FoldEnrichment**: reciprocal of RichFactor (1/RichFactor); **Zscore**: The *Z* score represents the standardized deviation of the observed enrichment from the expected enrichment under the null hypothesis; **Pvalue**: *P* value from hypergeometric (Fisher's exact test) for enrichment; **P.adjust**: Benjamini-Hochberg FDR *P* value correction; **Qvalue**: Storey's *Q* value method; **Count**: number of genes in experimental dataset background. **GeneID**: Affected genes within each GO term.

**Table S4. Ingenuity Pathway Analysis - enriched pathway terms.** Table of enriched pathways from Ingenuity Canonical Pathways analysis for CCI vs RAP at d14. Rows are Ingenuity Canonical Pathways, sorted by *Z* score. Column definitions: **Ingenuity Canonical Pathways**: Names of enriched pathways; **-log *P* value**: negative $\log_{10}$ of *P* value from Fisher's exact test (right-tailed); ***Z* score**: *Z* scores represent the standardized enrichment magnitude relative to expected background distribution (*Z* score ≥ 2 indicates activation, ≤ 2 indicates inhibition of the reported pathway based on IPA's functional annotations. **Molecules**: Affected genes within each pathway.

**Table S5. *P* values calculated from CPM under chromatin accessibility peaks.** (I) and (D), increased accessibility in the first named relative to second named condition, respectively; ns, not significant.

**Table S6. Patient characteristics analyzed by Omni-ATAC optimized for $CD66b^+$ MDSCs.** RAP, rapid recovery; CCI, chronic critical illness; HTN, hypertension; CHF, congestive heart failure; DM, diabetes mellitus; COPD, chronic obstructive pulmonary disease; GU, genitourinary; OSA, obstructive sleep apnea; AKI, acute kidney injury.
